# Tissue specificity-aware TWAS (TSA-TWAS) framework identifies novel associations with metabolic, immunologic, and virologic traits in HIV-positive adults

**DOI:** 10.1101/2020.08.31.273458

**Authors:** Binglan Li, Yogasudha Veturi, Anurag Verma, Yuki Bradford, Eric S. Daar, Roy M. Gulick, Sharon A. Riddler, Gregory K. Robbins, Jeffrey L. Lennox, David W. Haas, Marylyn D. Ritchie

**Affiliations:** Genomics and Computational Biology Program, University of Pennsylvania, Philadelphia, Pennsylvania, United States of America; Department of Genetics, University of Pennsylvania, Philadelphia, Pennsylvania, United States of America; Lundquist Institute at Harbor-UCLA Medical Center, Torrance, California, United States of America; Weill Cornell Medicine, New York, New York, New York, United States of America; University of Pittsburgh, Pittsburgh, Pennsylvania, United States of America; Harvard Medical School, Boston, Massachusetts, United States of America; Emory University School of Medicine, Atlanta, Georgia, United States of America; Departments of Medicine, Pharmacology, Pathology, Microbiology & Immunology, Vanderbilt University School of Medicine, Nashville, Tennessee, United States of America; Department of Internal Medicine, Meharry Medical College, Nashville, Tennessee, United States of America; Institute for Biomedical Informatics, University of Pennsylvania, Philadelphia, Pennsylvania, United States of America

## Abstract

As a type of relatively new methodology, the transcriptome-wide association study (TWAS) has gained interest due to capacity for gene-level association testing. However, the development of TWAS has outpaced statistical evaluation of TWAS gene prioritization performance. Current TWAS methods vary in underlying biological assumptions about tissue specificity of transcriptional regulatory mechanisms. In a previous study from our group, this may have affected whether TWAS methods better identified associations in single tissues versus multiple tissues. We therefore designed simulation analyses to examine how the interplay between particular TWAS methods and tissue specificity of gene expression affects power and type I error rates for gene prioritization. We found that cross-tissue identification of expression quantitative trait loci (eQTLs) improved TWAS power. Single-tissue TWAS (i.e., PrediXcan) had robust power to identify genes expressed in single tissues, but, had high false positive rates for genes that are expressed in multiple tissues. Cross-tissue TWAS (i.e., UTMOST) had overall equal or greater power and controlled type I error rates for genes expressed in multiple tissues. Based on these simulation results, we applied a tissue specificity-aware TWAS (TSA-TWAS) analytic framework to look for gene-based associations with pre-treatment laboratory values from AIDS Clinical Trial Group (ACTG) studies. We replicated several proof-of-concept transcriptionally regulated gene-trait associations, including *UGT1A1* (encoding bilirubin uridine diphosphate glucuronosyl transferase enzyme) and total bilirubin levels (p = 3.59×10^−12^), and *CETP* (cholesteryl ester transfer protein) with high-density lipoprotein cholesterol (p = 4.49×10^−12^). We also identified several novel genes associated with metabolic and virologic traits, as well as pleiotropic genes that linked plasma viral load, absolute basophil count, and/or triglyceride levels. By highlighting the advantages of different TWAS methods, our simulation study promotes a tissue specificity-aware TWAS analytic framework that revealed novel aspects of HIV-related traits.

publicly available.

## Introduction

Translating fundamental genetics research discoveries into clinical research and clinical practice is a challenge for biomedical studies of complex human traits [1,2]. Greater than 90% of complex trait-associated single-nucleotide polymorphisms (SNPs) identified via genome-wide association studies (GWAS) are located in noncoding regions of the human genome [3,4]. The difficulty in making connections between noncoding variants and downstream affected genes can hinder the translatability of GWAS discoveries to clinical research. The emerging transcriptome-wide association studies (TWAS) are a type of recently developed bioinformatics methodology that provide a means to address the challenge of GWAS translatability. TWAS mitigates the translational issue by integrating GWAS data with expression quantitative trait loci (eQTLs) information to perform gene-level association analyses. TWAS hypothesizes that SNPs act as eQTLs to collectively moderate the transcriptional activities of genes and thus influence complex traits of interest [5,6]. Accordingly, TWAS methods in general comprise two steps. The first step in TWAS is to impute the genetically regulated gene expression (GReX) for research samples in a tissue-specific manner. The second step is to conduct association analyses between GReX and the trait of interest to evaluate the gene-trait relationship for statistical significance [7-9]. Genome-wide eQTLs data are now available for various primary human tissues (e.g., liver, brain and heart) thanks to large-scale eQTL consortia including the Genotype-Tissue Expression (GTEx) project [10] and the eQTLGEN consortium [11]. The considerable centralized eQTL data have been fostering the development and application of TWAS.

While TWAS is an innovative and potentially powerful computational approach, several factors can influence TWAS. The choice of eQTL datasets matters for the performance of TWAS [12]. Most available eQTLs to date are identified in a tissue-by-tissue manner [5,10]. This approach, however, does not leverage the potential for shared transcriptional regulatory mechanisms across tissues, and can be limited by sample sizes of single tissues. One way to overcome this limitation is to take into consideration all available tissues, so as to increase sample sizes and improve the quality of eQTL datasets. We referred this type of eQTL detection method as the integrative tissue-based eQTL detection method [13-15]. Without a simulation study, however, it was unclear *how the choice of eQTL detection methods will impact TWAS*.

Another prominent question in TWAS studies is the choice of the association approaches. TWAS started with single-tissue association approaches, such as PrediXcan [5] and FUSION [6]. The most recent TWAS methods, such as UTMOST [15] and MulTiXcan [16], perform cross-tissue association analyses. Such TWAS methods evaluate whether a gene is significantly associated with a trait by integrating association data across tissues and adjusting for the statistical correlation structure among tissues. However, genes may vary substantially with regard to how they are regulated across tissues. When a gene is specifically expressed in a single or few tissues versus expressed in multiple tissues, *how will tissue specificity of gene expression affect TWAS power and type I error rates?*

Another appealing feature of TWAS is its capacity for tissue-specific association analyses thanks to the availability of tissue-specific eQTLs in a variety of primary human tissues. However, several recent studies revealed shared regulatory mechanisms across multiple human tissues [17] and showed that *cis*-eQTLs are less tissue-specific than other regulatory elements [10,11]. This suggests that TWAS can possibly identify genes in tissues that share biology with the causal tissue(s), but in fact are not the causal tissues for the trait of interest [18]. While TWAS is likely to identify false positive tissues, to date, the false positive rates of tissues in TWAS is unknown.

The above TWAS challenges can be summarized in two questions — *How does tissue specificity affect TWAS performance? How would this impact the choices of TWAS methods?* Available simulation strategies can be limited in answering these questions. Some have not taken into consideration the gene expression correlation structure across tissues [19,20]. Some assume a monogenic structure of transcriptional regulation [13-15,21], rather than the polygenic structure suggested by recent studies [10,22,23]. To address these issues, we applied a novel strategy to simulate eQTLs and gene expression of a wide range of tissue specificity (see **Methods**). We then applied different TWAS methods on the simulated datasets to assess power, type I error rates, and false positive rates of tissues. We found that the tissue specificity affected TWAS performance, with no single type of TWAS method being best for every type of genetic background of transcriptional regulation.

The simulation results motivated the development and implementation of an enhanced, tissue specificity-aware TWAS (TSA-TWAS) analytic framework. We tested the performance of TSA-TWAS analytic framework using AIDS Clinical Trials Group (ACTG) data (described in **Methods**). We showed that the TSA-TWAS was able to both replicate proof-of-concept gene-trait associations and identify novel trait-related genes. The simulation scheme highlighted the effects of tissue specificity on TWAS performance, and that TSA-TWAS could help better understand regulatory mechanisms that underlie complex human traits.

## Results

### Simulation design

We designed a novel simulation framework to investigate how the tissue specificities of eQTLs and gene expression affected TWAS power and type I error rates, and the choices of TWAS methods (Fig 1). We tested two representative eQTL detection methods, elastic net (implemented in PrediXcan [5]) and group LASSO (implemented in UTMOST [15]); and two gene-trait association approaches, Principal Component Regression (PC Regression; implemented in MulTiXcan [16]) and Generalized Berk-Jones test (GBJ test; implemented in UTMOST [15]) (Table 1).

**Fig 1.**
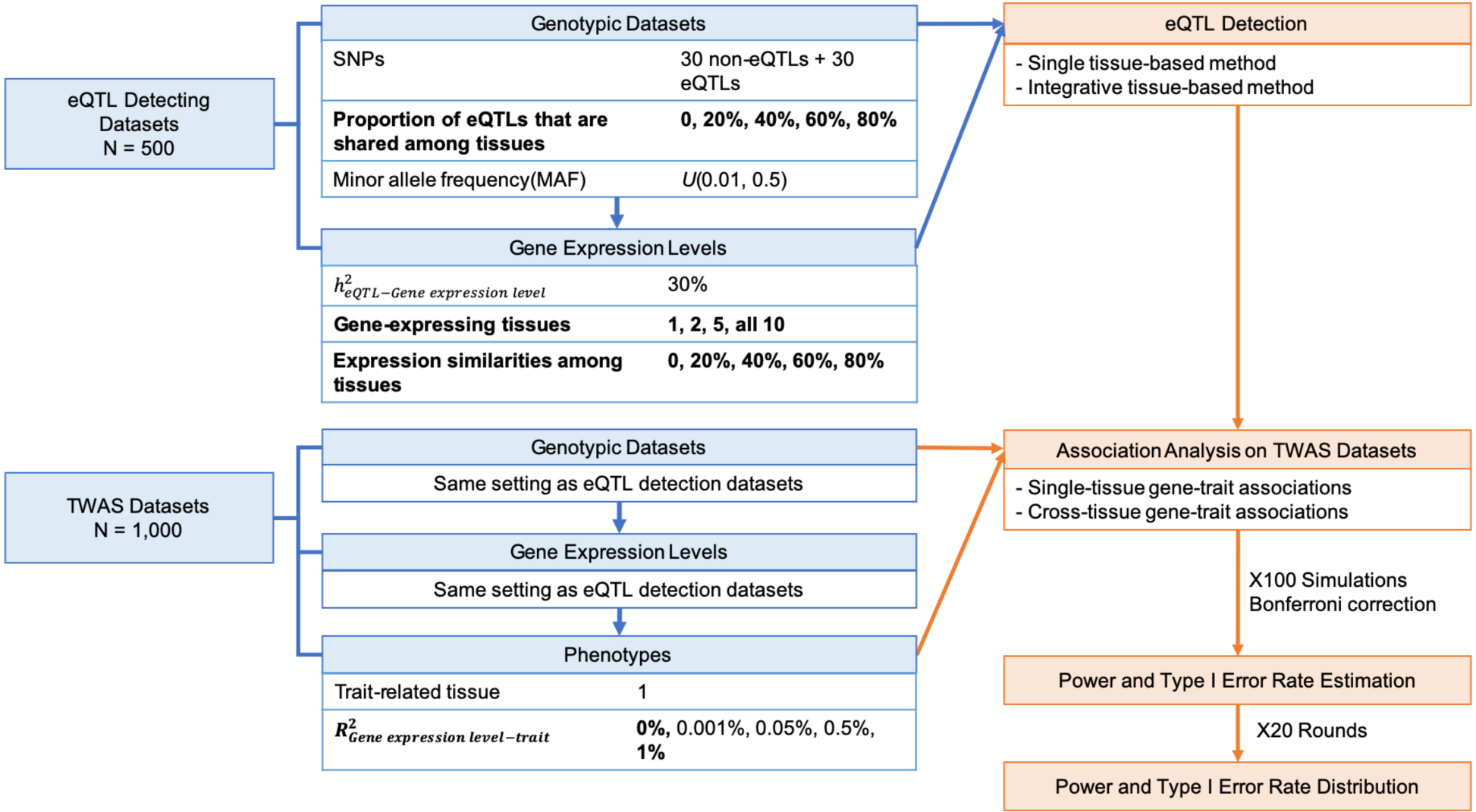
Cross-tissue TWAS simulation scheme. With the simulation parameters, we were able to generate SNP-gene-trait relations of varied tissue specificity backgrounds. In each replication, simulated datasets were divided into an eQTL detection dataset and a TWAS dataset. The former was used to identify eQTLs using different eQTL detection methods and the sample size was equivalent to that of GTEx. The detected eQTLs were then passed, separately, to the TWAS dataset to assist gene-level association tests. The TWAS dataset sample size was equivalent of that of the ACTG clinical trial dataset. Two types of gene-level association approaches estimated and ascribed p-values to the simulated gene-trait relations. In each replication, we simulated 100 different SNP-gene-trait pairs for one single point estimation of TWAS gene prioritization performance. All association p-values had been adjusted for the number of genes and tissues in each replication. 20 independent replications were conducted to obtain the distribution of TWAS performance statistics.

**Table 1.** TWAS methods tested in this simulation study.

| eQTL detection methods |  | Gene-trait association approaches |  | Equivalent developed TWAS methods | PMID |
| --- | --- | --- | --- | --- | --- |
| Type | Name | Type | Name |  |  |
| Single tissue-based | Elastic net | Single-tissue association | Linear or logistic regression | PrediXcan | 31086352 |
| Integrative tissue-based | Group LASSO | Single-tissue association | Linear or logistic regression | Single-tissue UTMOST | 30804563 |
| Single tissue-based | Elastic net | Cross-tissue association | Principal component regression | MultiXcan | 30668570 |
| Integrative tissue-based | Group LASSO | Cross-tissue association | Generalized Berk-Jones test | Cross-tissue UTMOST | 30804563 |

Tissue-specific eQTLs were defined as those that were only functioning in one single tissue. Multi-tissue eQTLs were defined as those that had regulatory effect across all gene-expressing tissues (see **Methods**). We generated genes that had different genetic makeup of tissue-specific and multi-tissue eQTLs in a gene to evaluate the influence of tissue specificity of eQTLs on TWAS performance.

Tissue specificity of gene expression was determined by the number of gene-expressing tissues and the similarity of gene expression levels across tissues. (see **Methods**). Tissue-specific genes were those specifically expressed in only one or two tissues. Ubiquitously expressed genes were those expressed in all ten simulated tissues with high gene expression similarity (expression similarity = 60%, 80%). Differentially expressed and similarly expressed genes were those having distinctive gene expression levels (gene expression similarity = 0, 20% 40%) or highly correlated gene expression levels across tissues (gene expression similarity = 60%, 80%), respectively, regardless of the number of gene-expressing tissues. To evaluate the impact of tissue specificity of gene expression on TWAS performance, we generated genes that were expressed in varied numbers of tissues and had diverse gene expression similarities across tissues.

In addition, we designed different strength of gene-trait associations defined by 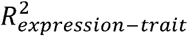 (the proportion of phenotypic variation explained by gene expression levels), but the reported results by default were the cases under 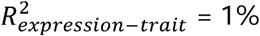. Only continuous traits were evaluated in this simulation study, in accordance with ACTG baseline laboratory values in the real-world application dataset.

### Power of different TWAS methods

We did not observe any obvious effect of tissue-specificity of eQTLs on TWAS power, except for ubiquitously expressed genes. TWAS, specifically group LASSO (implemented in UTMOST[15]), had greater power to prioritize ubiquitously expressed genes that were mostly regulated by multi-tissue eQTLs than those that were not (Fig 2, bottom row).

**Fig 2.**
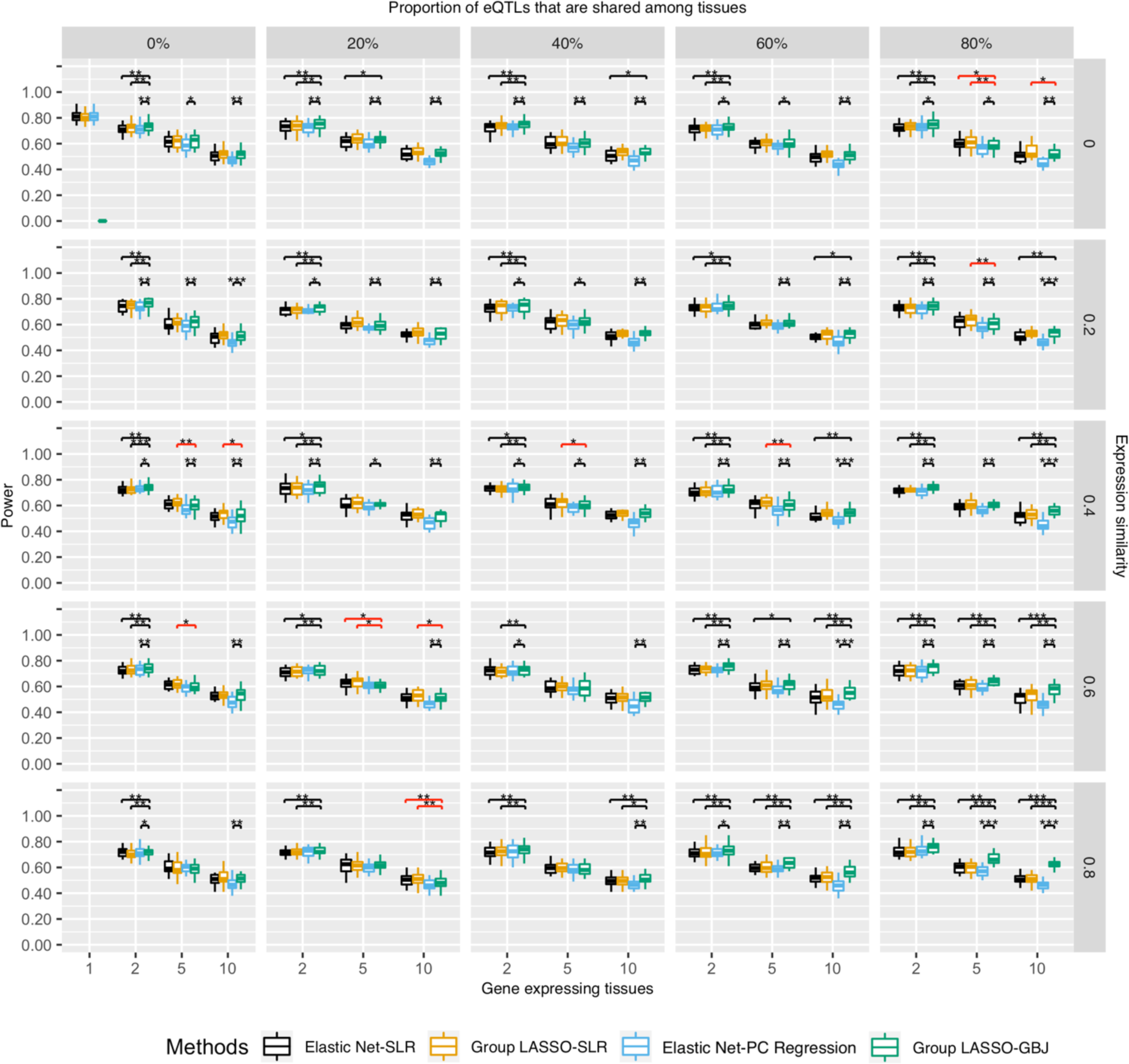
Power of different TWAS methods in prioritizing genes of varied tissue specificity properties. Power was the proportion of successfully identified gene-trait associations in the causal tissue out of all simulations. X-axis is the number of gene-expressing tissues. Each column stands for the proportion of eQTLs that are shared among tissues for a gene. Each row is the similarity of gene expression profiles across tissues which is estimated by correlation. Moving from the top left to the bottom right is a gradient spectrum from tissue-specific genes to broadly expressed genes. The colors represent different TWAS methods and y-axis is the power. For tissue-specific genes at the top left, single-tissue TWAS (Elastic Net-SLR) and cross-tissue TWAS (Group LASSO-GBJ) had similar power. For broadly expressed genes at the bottom right, cross-tissue TWAS (Group LASSO-GBJ) had greater power. Brackets showed pairwise comparison of power between the Group LASSO-GBJ and other TWAS methods using Wilcoxon Signed-rank Test. Black brackets were cases where Group LASSO-GBJ had higher power than other three methods; red brackets were cases where Group LASSO-GBJ had lower power than other three methods. *p-value < 0.05, **p-value < 0.01, ***p-value < 0.0001.

We then asked how eQTL detection methods affected TWAS gene-prioritization power, and whether one eQTL detection method was preferred over another. We found that the integrative tissue-based eQTL detection method had, on average, approximately 2% greater power than the single-tissue method. Take differentially expressed genes for instance, eQTLs identified via the Group LASSO led to 53.8% gene prioritization power of TWAS and eQTLs identified via the Elastic Net led to 50.7% power (Wilcoxon Signed-rank Test p = 5.85×10^−4^; S4 Fig, top right corner). More pairwise comparison results among all TWAS methods can be found in S1 Table. Overall, TWAS gained slightly more power when using eQTLs identified in an integrative tissue context.

Gene-trait association approaches affected TWAS power more so than did choice of eQTL detection method. For tissue-specific genes or differentially expressed genes, SLR consistently had equal or greater power (average 70%) than the cross-tissue association approaches (PC regression and GBJ test; Fig 2, top left triangle). For ubiquitously expressed genes or similarly expressed genes, GBJ test had equal or greater power than the single-tissue association approach (SLR; Fig 2, bottom right triangle). Especially for ubiquitously expressed genes, GBJ test had statistically significant greater power (62%) compared to SLR (51%) (Fig 2, bottom right corner, Wilcoxon Signed-rank Test p = 9.4×10^−5^).

The group LASSO-GBJ test (implemented in UTMOST) had a greater power to prioritize similarly or ubiquitously expressed genes. For genes that were expressed in five tissues, power of the group LASSO-GBJ test increased from 62.2% for differentially expressed genes (Fig 2, top left corner) to 66.6% for similarly expressed gene (Fig 2, bottom right corner). For genes that were expressed in all ten tissues, power of the group LASSO-GBJ test increased from 51.2% for differentially expressed genes (Fig 2, top left corner) to 61.9% for ubiquitously expressed gene (Fig 2, bottom right corner). Moreover, the group LASSO-GBJ test showed equal or statistically significant greater power than other TWAS methods in 65 of the 76 simulated scenarios (∼84%). Black brackets in Fig 2 showed cases where Group LASSO-GBJ had higher power than other three methods; red brackets showed cases where Group LASSO-GBJ had lower power than other three methods. Comprehensive statistical test results of power differences are available in S4 Fig and S1 Table. However, GBJ test cannot handle the case where the gene was only expressed in one single tissue.

Overall, the group LASSO-GBJ test had equal or greater power in prioritizing genes that were expressed in multiple tissues. Single-tissue association approaches (e.g. SLR) had greater power and robust performance in prioritizing tissue-specific genes.

The strength of gene-trait associations affected TWAS gene prioritization power. The stronger the gene-trait associations, the greater the power for TWAS gene prioritization (Fig 2, S5-7 Figs).

### Type I error rates of various TWAS methods

All TWAS methods had well-controlled type I error rates (≤ 5%; Fig 3, S2 Table). Significance thresholds in this simulation were corrected using the Bonferroni approach to control for family-wise error rate. All single-tissue association approaches (Elastic Net-SLR and Group LASSO-SLR) had less type I error rates than the cross-tissue associations approaches (Wilcoxon Signed-rank Test p < 0.01, S8 Fig). Both GBJ test and PC regression had average type I error rates of approximately 5%. The GBJ test showed statistically significant lower type I error rates than PC regression for ubiquitously expressed genes (Wilcoxon Signed-rank p < 0.05, S8 Fig, S2 Table).

**Fig 3.**
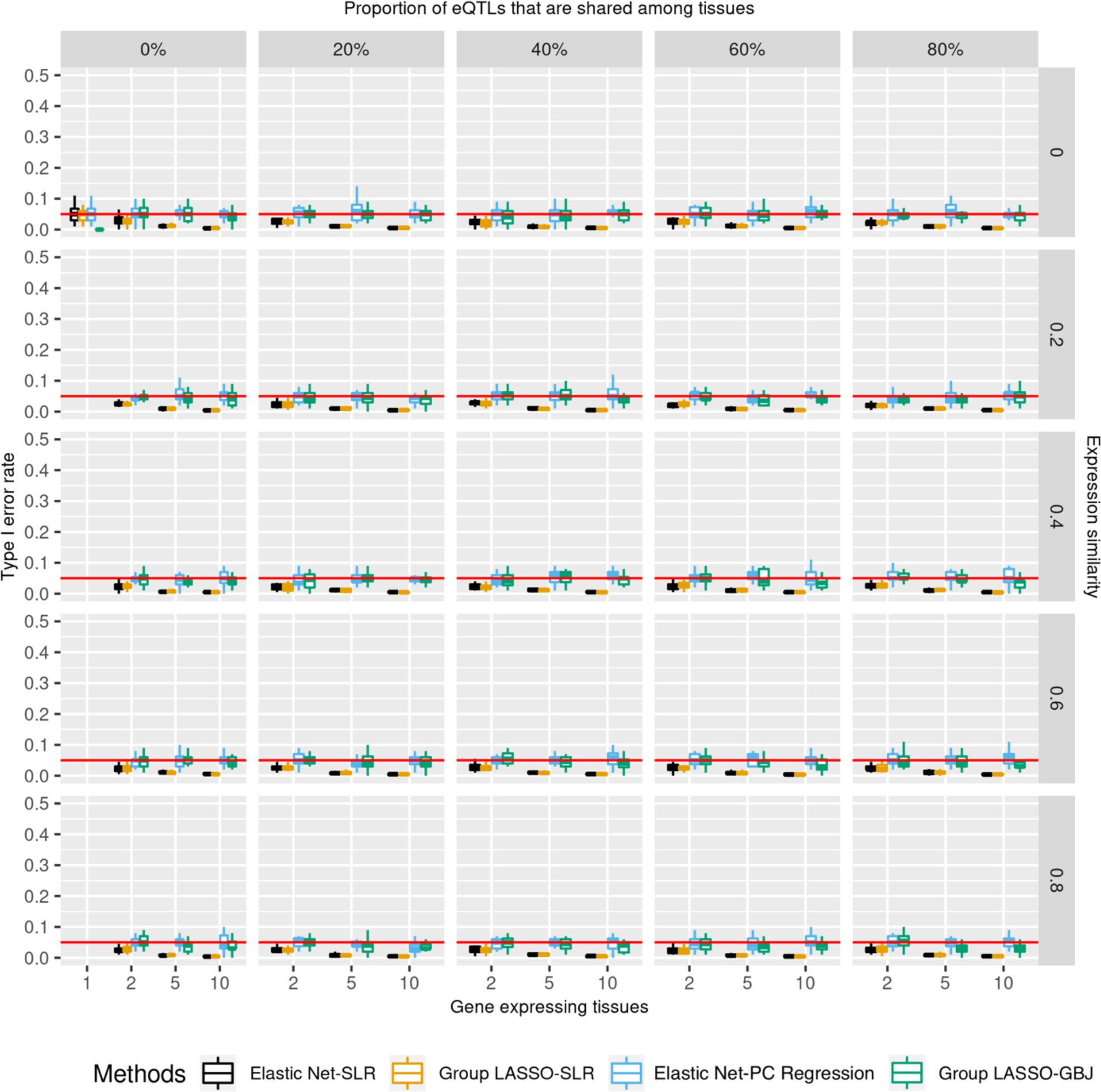
Type I error rates of different TWAS methods in prioritizing genes of diverse tissue specificity properties. Type I error rate was the probability that TWAS wrongly identified a gene-trait association as significant while there was not any signal simulated in the dataset. Association p-values were controlled for the number of genes and tested tissues. X-axis is the number of gene-expressing tissues. Each column stands for the proportion of eQTLs that are shared among tissues for a gene. Each row is the similarity of gene expression profiles across tissues which is estimated by correlation. Moving from the top left to the bottom right is a gradient spectrum from tissue-specific genes to broadly expressed genes. The colors represent different TWAS methods and y-axis is the type I error rate. All TWAS methods had controlled type I error rates (≤5%).

### False positives of statistically significant tissues

If not corrected for the number of tested tissues, single-tissue TWAS would have greater power (S9 Fig), but also a higher false positive rate for tissues (S10 Fig). False positive rates of tissues were at least 10% for genes that were expressed in more than one tissue. In effect, while the genes might be related to a trait of interest, 10% of statistically significant results pointed to wrong tissues. The false positive rate of tissues proportionally increased with the number of gene-expressing tissues. The highest false positive rates were seen in the case of ubiquitously expressed genes (S10 Fig, bottom right corner), which on average, had an 84% false positive rate based on 20 random replications. This suggested that any single-tissue TWAS may have 10-84% false positive rate tissues associations if not adjusted for the number of tested tissues.

Adjusting for the number of tested tissues reduced the false positive rates somewhat, but number-wise, the false positive rate may remain quite high. False positive rates of tissues were relatively controlled at approximately 5% for tissue-specific genes (Fig 4, top left corner). False positive rates still increased with the number of tissues in which a gene was expressed (Fig 4). Genes expressed in ten tissues had at least on average a 24% false positive rate. False positive rates were as high as 77% for ubiquitously expressed genes (Fig 4, bottom right corner).

**Fig 4.**
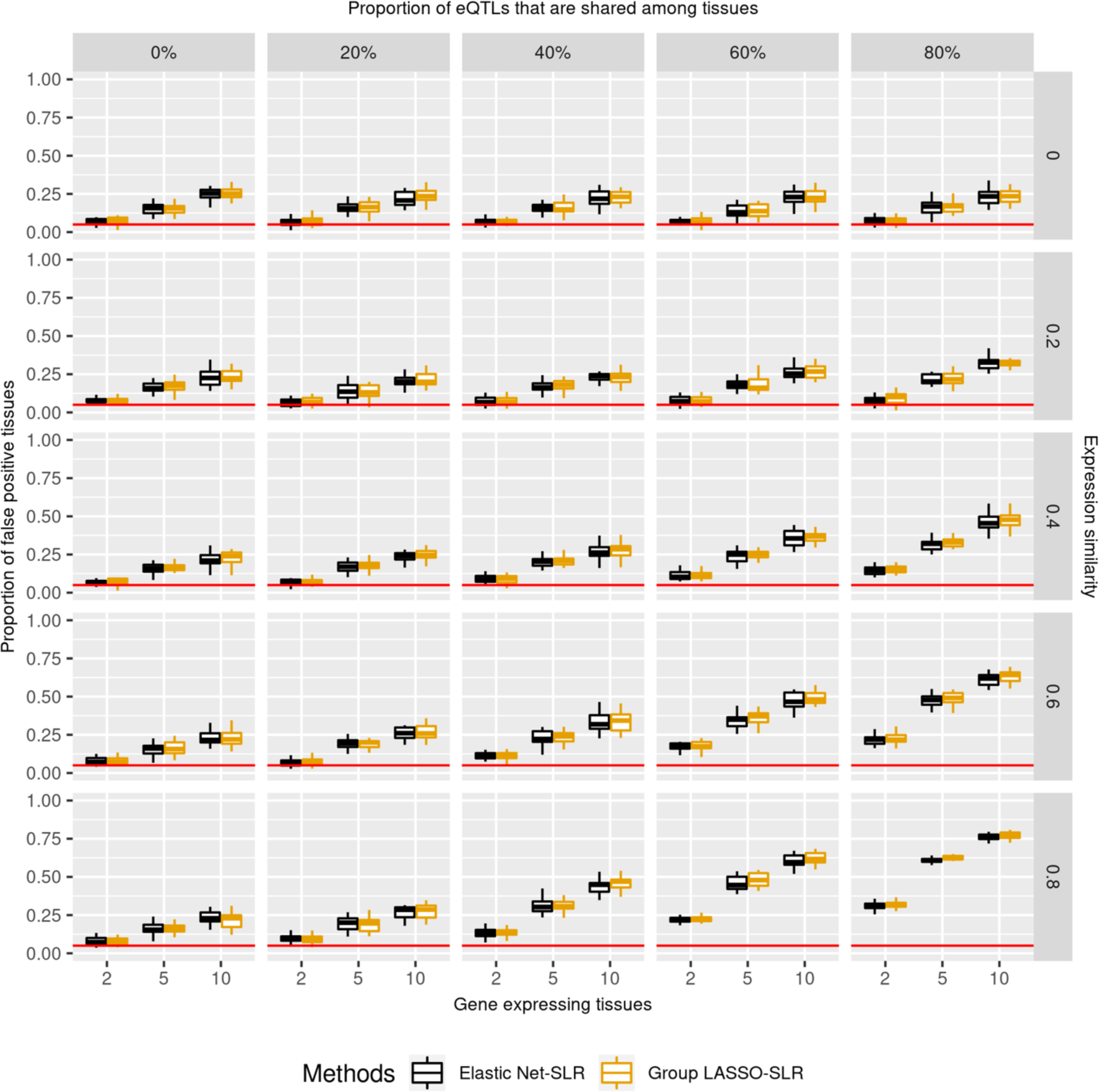
False positive rates of tissues among statistically significant results. False positive rates were the proportion of significant associations found in trait-irrelevant tissues amongst all significant results. Association p-values were controlled for the number of genes and tested tissues. X-axis is the number of gene-expressing tissues. Each column stands for the proportion of eQTLs that are shared among tissues for a gene. Each row is the similarity of gene expression profiles across tissues which is estimated by correlation. Moving from the top left to the bottom right is a gradient spectrum from tissue-specific genes to broadly expressed genes. Colors represent different TWAS methods and y-axis is the false positive rate of tissues among statistically significant results. Single-tissue TWAS wrongly identified 5% and 77% trait-irrelevant tissues for tissue-specific and broadly expressed genes, respectively.

### Validation and support of simulation design

To evaluate whether our simulation findings would translate from in silico parameter designs to real world scenarios, we designed a Monte Carlo simulation process to estimate the trait heritability behind various genetic scenarios (S11 Fig). The results suggested that 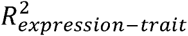 increased with trait heritability (S12 Fig). Heritability of traits with 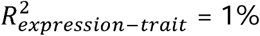 were estimated to be on average 1% (standard error (s.e.) = 0.059%) which were derived from multiple, repeated random sampling. In contrast, the minor allele frequencies (MAF) of eQTLs had almost no effect on trait heritability. This suggested that trait heritability positively influenced the strength of gene-trait associations in TWAS. In other words, if a trait was moderated by genetic factors through differential gene expression, the greater a trait’s heritability is, the stronger the associations were in TWAS.

### Designing the TSA-TWAS analytic framework

Our simulation suggested an influence of tissue specificity on TWAS performance. Thus, we designed a TSA-TWAS analytic framework to balance trade-offs among power, type I error rates, and false positive rates of tissues and to take into consideration the distribution of GReX (S13 Fig). The idea was illustrated in Fig 5. When trait-related tissue(s) are known, we recommend single-tissue TWAS in the known related tissues only. Additionally, we recommend using eQTLs identified by integrative tissue-based eQTL detection methods (for example, group LASSO), which showed slightly greater power. In contrast, if trait-related tissue(s) are uncertain, it may be better to stratify genes based on the number of tissues in which the genes are predicted to be expressed. For genes predicted to be expressed in just one tissue, single-tissue TWAS will have greater power and can provide information on trait-related tissues. For genes that are expressed multiple tissues, cross-tissue TWAS will provide overall equal or greater power, as well as controlled type I error rates.

**Fig 5.**
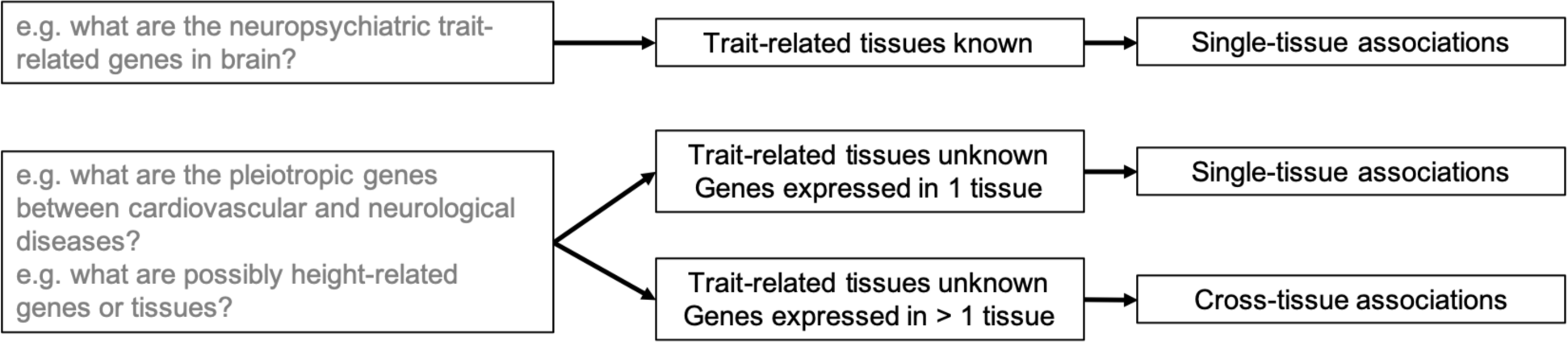
A proposed TSA-TWAS analytic framework that leverages TWAS performance on genes of different tissue specificity properties. The framework proposed based on our simulations is as follows: If trait-related tissue(s) are known for a trait or disease of interest, run single-tissue TWAS, for example, PrediXcan. If trait-related tissue(s) are unknown, run cross-tissue TWAS (UTMOST) on the genes that are expressed in more than one tissue and run single-tissue TWAS (PrediXcan) on the genes that are expressed in one single tissue.

**Fig 6.**
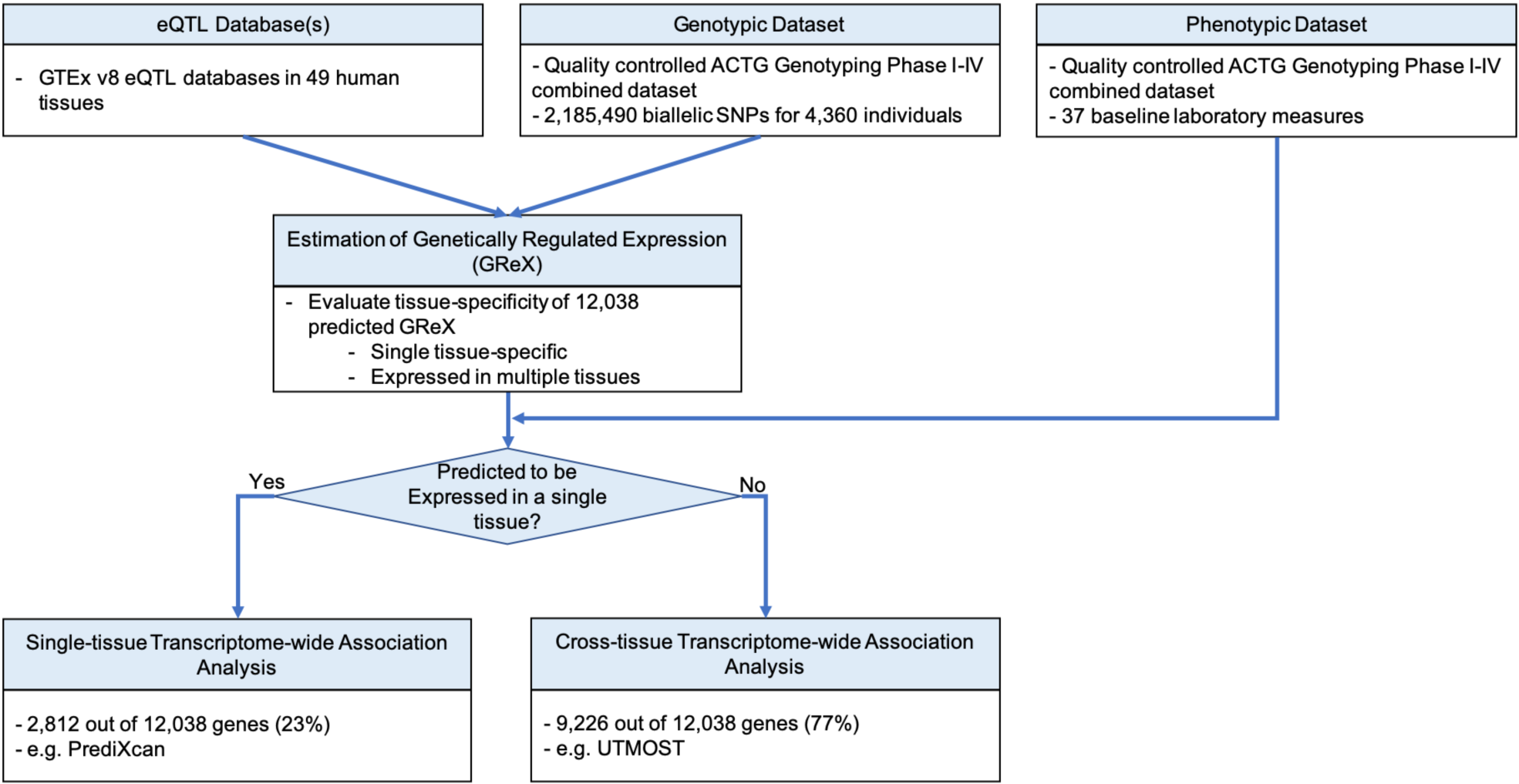
The TSA-TWAS analytic framework for the ACTG combined genotyping phase I-IV baseline laboratory traits. Approximately 2.2 million SNPs, 4,360 individuals, and 37 baseline laboratory traits survived the QC. UTMOST eQTL models were used to impute GReX of a total of 12,038 genes in 49 tissues. 2,812 genes (23%) had GReX in one single tissue, and 9,226 genes (77%) had GReX in more than one tissue.

### TSA-TWAS replicated known associations

We applied TSA-TWAS to 37 baseline laboratory values from a combined dataset of five clinical trials from AIDS Clinical Trials Group (ACTG) with available genotype data (N = 4,360; Table 2). We first imputed the GReX to distinguish genes whose GReX were only expressed in one tissue versus multiple tissues. Genes expressed in just one tissue comprised 2,812 (23%) of 12,038 genes on which data were available. The remaining 9,226 (77%) genes had GReX in multiple tissues. Genes expressed in one, and in more than one tissue were tested for associations with baseline laboratory values using single-tissue, and by cross-tissue gene-trait association approaches, respectively (see **Methods**). TSA-TWAS found in total 83 statistically significant gene-trait associations, comprising 45 distinct genes and 10 traits (Fig 7).

**Table 2.** Summary statistics of the ACTG genotyping phase I-IV baseline laboratories.

| Trait | Sample Size | Mean | Std. Dev. | Min | Max | Transformation | Unit | Description |
| --- | --- | --- | --- | --- | --- | --- | --- | --- |
| Albumin | 1216 | 4.05 | 0.44 | 1.80 | 5.30 |  | g/dL | week 0 albumin (Alb, g/dL) |
| Bicarbonate | 3971 | 26.01 | 2.94 | 12.00 | 35.00 |  | mmol/L | week 0 bicarbonate (Bicarb, mmol/L) |
| Calcium | 1336 | 9.17 | 0.44 | 7.40 | 10.80 |  | mg/dL | week 0 calcium (Ca, mg/dL) |
| Chloride | 4048 | 103.27 | 2.94 | 88.00 | 117.00 |  | mmol/L | week 0 chloride (Cl, mmol/L) |
| Cholesterol | 4286 | 159.27 | 36.80 | 5.90 | 414.00 |  | mg/dL | week 0 cholesterol (Chol, mg/dL) |
| Creatinine | 4100 | 0.91 | 0.20 | 0.05 | 2.80 |  | mg/dL | week 0 creatinine (Creat, mg/dL) |
| HDL-c | 2376 | 37.31 | 12.78 | 3.90 | 148.00 |  | mg/dL | week 0 HDL-c (HDL-c, mg/dL) |
| Hemoglobin | 4293 | 13.49 | 1.77 | 6.00 | 20.20 |  | g/dL | week 0 hemoglobin (Hgb, g/dL) |
| Absolute basophil count | 2526 | 1.44 | 0.32 | 0.00 | 3.39 | Log<sub>10</sub> | cells/mm<sup>3</sup> | log<sub>10</sub> transformed week 0 absolute basophil count (Baso, cells/mm<sup>3</sup>) |
| Absolute eosinophil count | 3932 | 2.06 | 0.40 | 0.18 | 3.55 | Log<sub>10</sub> | cells/mm<sup>3</sup> | log<sub>10</sub> transformed week 0 absolute eosinophil count (Eos, cells/mm<sup>3</sup>) |
| Alkaline phosphatase | 4226 | 1.88 | 0.15 | 0.70 | 2.72 | Log<sub>10</sub> | U/L | log<sub>10</sub> transformed week 0 alkaline phosphatase (AlkP, U/L) |
| ALT | 4233 | 1.48 | 0.27 | 0.04 | 2.81 | Log<sub>10</sub> | U/L | log<sub>10</sub> transformed week 0 ALT (ALT, U/L) |
| Absolute lymphocyte count | 4149 | 3.11 | 0.24 | 0.92 | 4.03 | Log<sub>10</sub> | cells/mm<sup>3</sup> | log<sub>10</sub> transformed week 0 absolute lymphocyte count (Lymph, cells/mm<sup>3</sup>) |
| Absolute monocyte count | 4116 | 2.58 | 0.21 | 0.66 | 3.69 | Log<sub>10</sub> | cells/mm<sup>3</sup> | log<sub>10</sub> transformed week 0 absolute monocyte count (Mono, cells/mm<sup>3</sup>) |
| Amylase | 1026 | 1.85 | 0.20 | 1.11 | 2.89 | Log<sub>10</sub> | U/L | log<sub>10</sub> transformed week 0 amylase (Amyl, U/L) |
| Absolute neutrophil count | 4277 | 3.32 | 0.21 | 2.28 | 4.67 | Log<sub>10</sub> | cells/mm<sup>3</sup> | log<sub>10</sub> transformed week 0 absolute neutrophil count (ANC, cells/mm<sup>3</sup>) |
| AST | 4235 | 1.49 | 0.21 | 0.48 | 2.81 | Log<sub>10</sub> | U/L | log<sub>10</sub> transformed week 0 AST (AST, U/L) |
| BUN | 4221 | 1.08 | 0.15 | -0.22 | 2.17 | Log<sub>10</sub> | mg/dL | log<sub>10</sub> transformed week 0 BUN (BUN, mg/dL) |
| CK | 1360 | 1.97 | 0.38 | -0.05 | 3.79 | Log<sub>10</sub> | U/L | log<sub>10</sub> transformed week 0 CK (CK, U/L) |
| Fasting glucose | 3233 | 1.93 | 0.08 | 1.52 | 2.64 | Log<sub>10</sub> | mg/dL | log<sub>10</sub> transformed week 0 fasting glucose (Gluc fasting, mg/dL) |
| Glucose (Log<sub>10</sub>) | 3031 | 1.93 | 0.08 | 1.70 | 2.77 | Log<sub>10</sub> | mg/dL | log<sub>10</sub> transformed week 0 glucose (Gluc, mg/dL) |
| LDL-c | 3539 | 1.95 | 0.16 | 0.00 | 2.57 | Log<sub>10</sub> | mg/dL | log<sub>10</sub> transformed week 0 LDL-c (LDL-c, mg/dL) |
| Lipoprotein | 1118 | 1.58 | 0.32 | 0.30 | 2.85 | Log<sub>10</sub> |  | log<sub>10</sub> transformed week 0 lipoprotein |
| Platelet count | 4263 | 2.30 | 0.15 | 1.15 | 3.34 | Log<sub>10</sub> | x10E9/L | log<sub>10</sub> transformed week 0 platelet count (Plat, x10E9/L) |
| Total bilirubin | 4202 | -0.31 | 0.21 | -1.00 | 0.49 | Log<sub>10</sub> | mg/dL | log<sub>10</sub> transformed week 0 total bilirubin (TBili, mg/dL) |
| Triglyceride | 4318 | 2.07 | 0.25 | 1.08 | 3.45 | Log<sub>10</sub> | mg/dL | log<sub>10</sub> transformed week 0 triglyceride (Trig, mg/dL) |
| White blood cell count | 4279 | 0.62 | 0.16 | -0.05 | 1.49 | Log<sub>10</sub> | x10E3 cells/cu | log<sub>10</sub> transformed week 0 white blood cell count (WBC, x10E3) |
|  |  |  |  |  |  |  | mm | cells/cu mm) |
| Hematocrit | 4274 | 39.83 | 5.10 | 1.00 | 62.10 |  | percent | week 0 hematocrit (Hct, percent) |
| Phosphate | 3261 | 3.44 | 0.61 | 0.80 | 7.70 |  | mg/dL | week 0 phosphate (Phos, mg/dL) |
| Potassium | 4062 | 4.15 | 0.39 | 2.00 | 8.00 |  | mmol/L | week 0 potassium (K, mmol/L) |
| Sodium | 4067 | 138.88 | 2.80 | 123.00 | 151.00 |  | mmol/L | week 0 sodium (Na, mmol/L) |
| CD4 count | 4358 | 14.78 | 6.46 | 0.00 | 36.55 | Square root | cells/mm<sup>3</sup> | square root of absolute CD4 count at week 0 |
| Viral load | 4358 | 4.75 | 0.72 | 2.02 | 7.11 | Log<sub>10</sub> | copies/dL | week 0 viral load RNA |
| Fasting cholesterol | 4136 | 158.42 | 36.24 | 6.10 | 414 |  | mg/dL | week 0 fasting cholesterol |
| Fasting HDL-c | 4126 | 1.56 | 0.15 | 0.60 | 2.20 | Log<sub>10</sub> | mg/dL | log<sub>10</sub> transformed week 0 fasting HDL-c |
| Fasting LDL-c | 4042 | 1.95 | 0.15 | 0.85 | 2.57 | Log<sub>10</sub> | mg/dL | log<sub>10</sub> transformed week 0 fasting LDL-c |
| Fasting triglyceride | 3888 | 2.05 | 0.24 | 1.08 | 2.45 | Log<sub>10</sub> | mg/dL | log<sub>10</sub> transformed week 0 fasting triglycerides |

**Fig 7.**
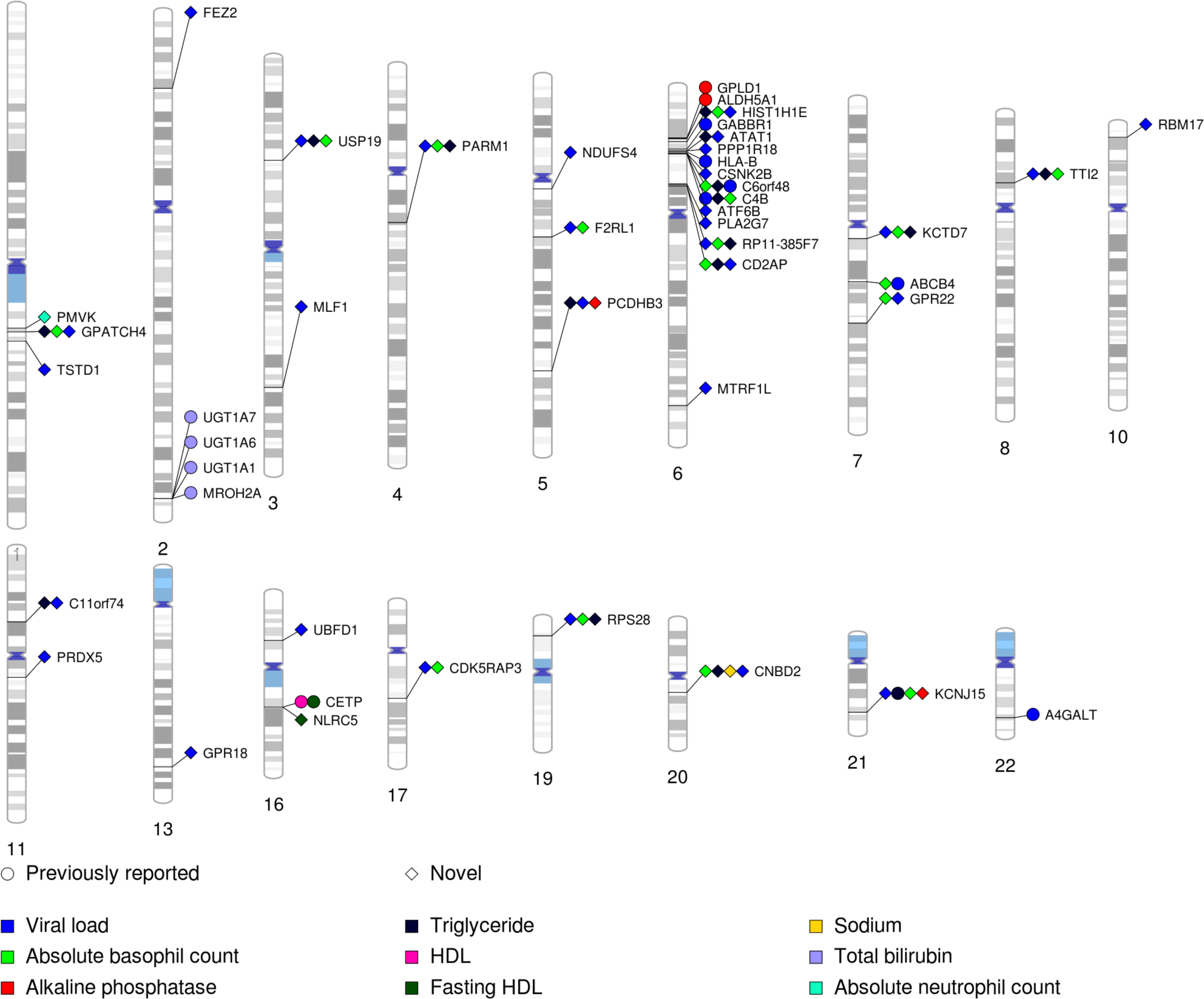
PhenoGram of statistically significant gene-trait associations identified by the TSA-TWAS analytic framework. We plotted the associations with p-value < 1.12 × 10^−7^. Each association is arranged according to the SNP location on each chromosome and the points are color-coded by baseline laboratory values. Diamonds represented previously reported or replicated associations, and circle represented novel associations identified in this study.

TSA-TWAS replicated several previously reported risk genes for certain baseline lab values (Table 3). The lowest p-values for association were observed between total plasma bilirubin levels and several genes on chromosome 2, nearby or overlapping *UGT1A1*. These included *MROH2A* (p = 1.39×10^−12^), which has been previously reported by GWAS of various populations [24-27], *UGT1A6* (p = 2.78×10^−15^), *UGT1A7* (p = 4.51×10^−12^) and *UGT1A1* (p = 3.59×10^−12^) [24,25,27,28]. We replicated the well-known association between *CETP* and high-density lipid-cholesterol levels (HDL-c; p = 4.49×10^−12^) [29]. Association was also found between *GPLD1* and plasma alkaline phosphatase levels (p = 1.08×10^−11^) [30]. *GPLD1* encodes a glycosylphosphatidylinositol-degrading enzyme that releases attached proteins from the plasma membrane and engages in regulation of alkaline phosphate activities. Other replicated discoveries included association between *ALDH5A1* and plasma alkaline phosphatase levels (p = 1.79×10^−11^) [31], *C6orf48* and absolute basophil count (p = 1.69×10^−12^) [32], *KCNJ15* and plasma triglyceride levels (p = 3.18×10^−1^) [33].

**Table 3.** Replicated associations related to HIV baseline laboratory values identified by TSA-TWAS.

| Trait | Gene | Chromosome | TSS | P |
| --- | --- | --- | --- | --- |
| Alkaline phosphatase | *GPLD1* | 6 | 24428177 | 1.08E-11 |
|  | *ALDH5A1* | 6 | 24494852 | 1.79E-11 |
| Fasting HDL | *CETP* | 16 | 56961850 | 4.49E-12 |
| HDL | *CETP* | 16 | 56961850 | 4.49E-12 |
| Total bilirubin | *UGT1A6* | 2 | 233692866 | 2.78E-15 |
|  | *MROH2A* | 2 | 233775679 | 1.39E-12 |
|  | *UGT1A1* | 2 | 233760248 | 3.59E-12 |
|  | *UGT1A7* | 2 | 233681938 | 4.51E-12 |
| Triglyceride | *KCNJ15* | 21 | 38256698 | 3.18E-13 |
| Viral load | *C4B* | 6 | 32014762 | 4.11E-15 |
|  | *GABBR1* | 6 | 29602228 | 1.14E-12 |
|  | *ABCB4* | 7 | 87401697 | 1.07E-11 |
|  | *HLA-B* | 6 | 31269491 | 1.15E-11 |
|  | *C6orf48* | 6 | 31834608 | 2.32E-11 |
|  | *A4GALT* | 22 | 42692121 | 8.39E-11 |

We have additionally replicated several genes’ association with plasma viral loads in HIV-positive adults, including *A4GALT* (p = 8.39×10^−11^) [34], *ABCB4* (p = 1.07×10^−11^) [35], *C4B* (p = 4.11×10^−15^) [36], *GABBR1*(p = 1.14×10^−12^) [37], and HLA-B (p = 1.15×10^−11^) [38].

### Novel genes prioritized by the TSA-TWAS

In addition to the above replications, TSA-TWAS identified novel associations with plasma viral load (Table 4). For instance, *PRDX5* (p = 7.01×10^−14^, which encodes a member of the peroxiredoxin family of antioxidant enzymes) was associated with plasma viral load with great significance. Several novel genes were first time reported to be associated with certain baseline laboratory values, which were otherwise associated with other traits by previous studies. For instance, *ATF6B* is a protein-coding gene that encodes a transcription factor in the unfolded protein response (UPR) pathway during ER stress and it has been associated with HIV-associated neurocognitive disorders in previous research. In our study, ATF6B associates with plasma viral load (p = 2.83×10^−9^).

**Table 4.**
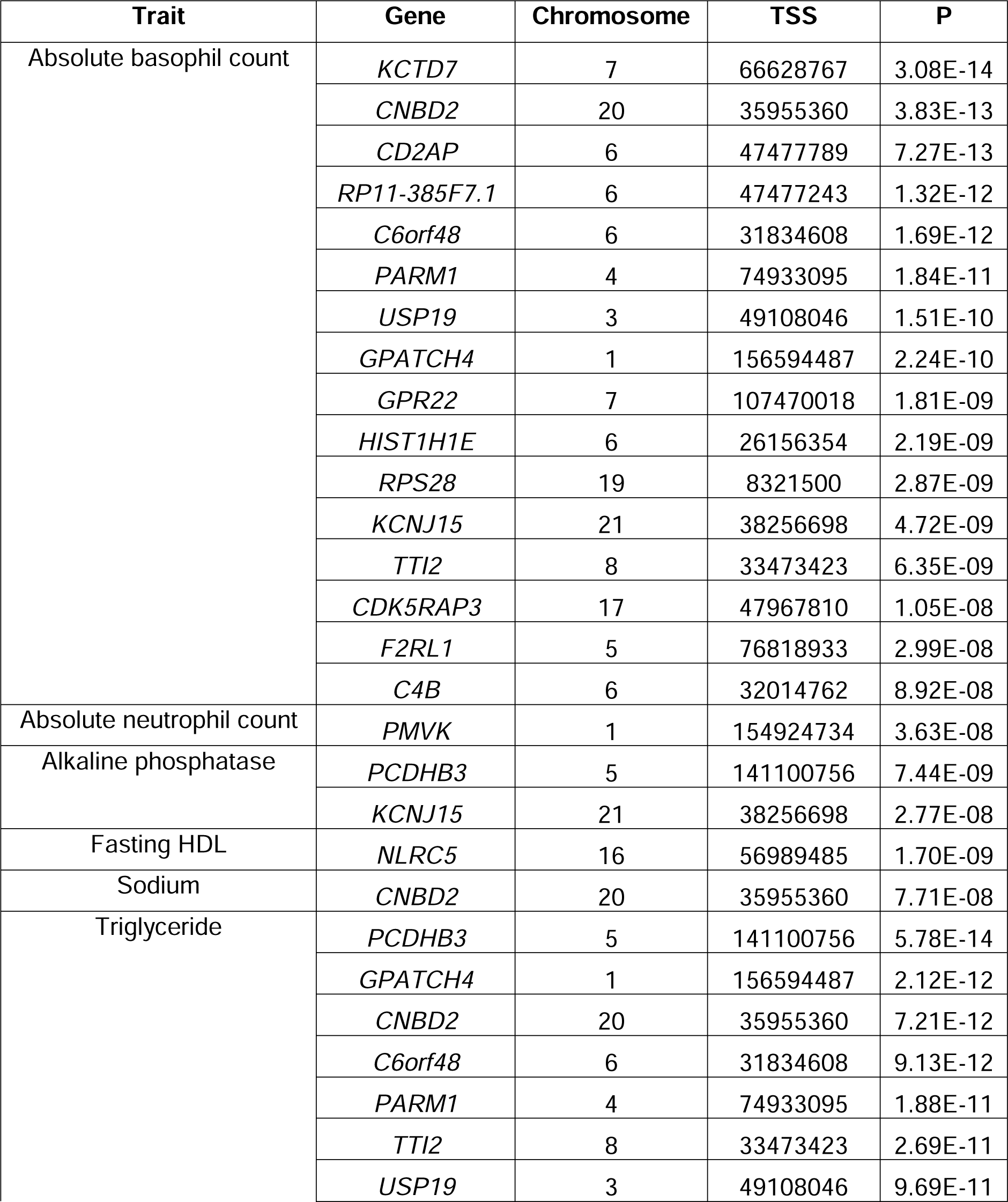
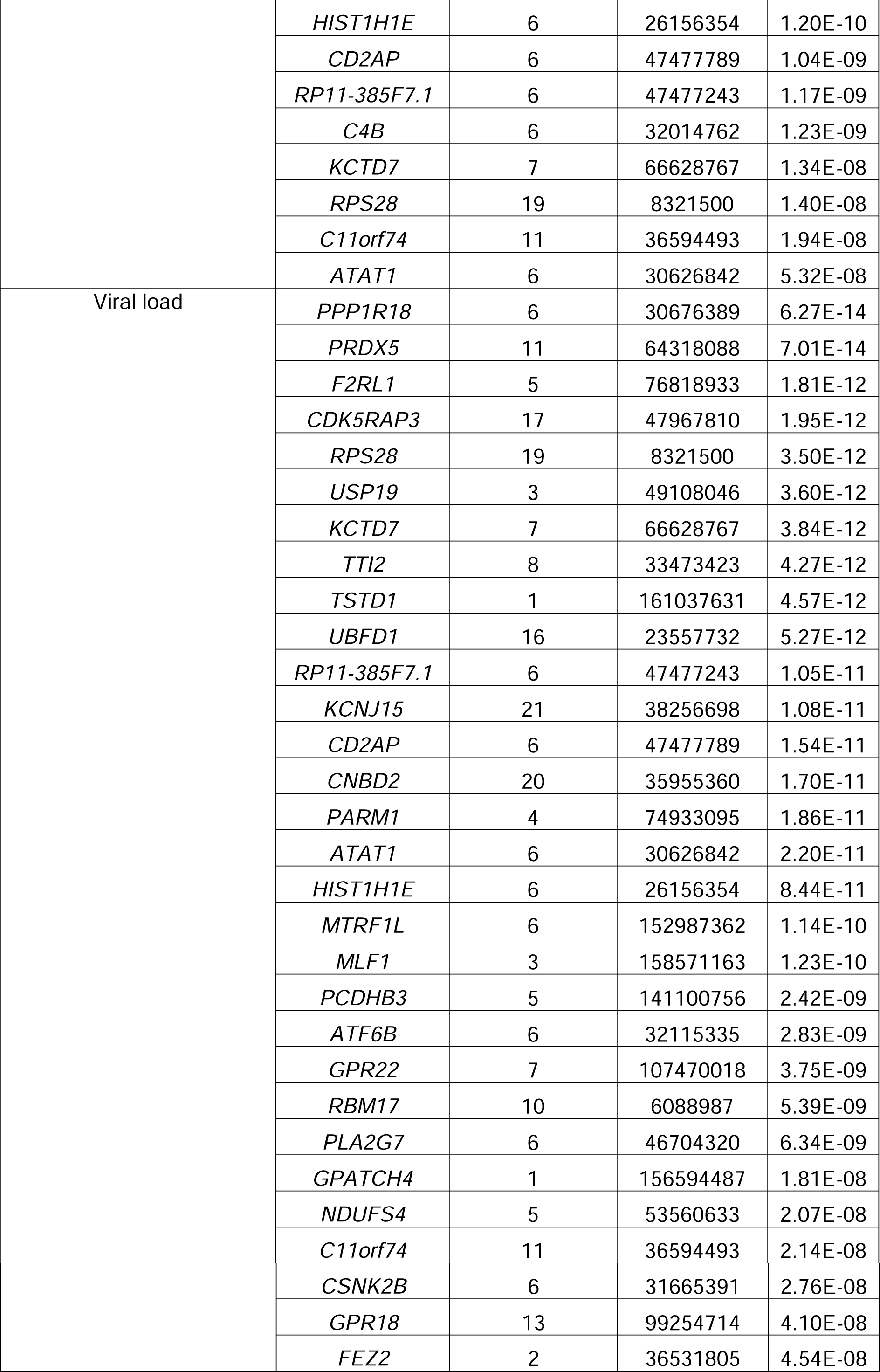
Novel associations related to HIV baseline laboratory values identified by TSA-TWAS.

### Pleiotropic genes associated with baseline laboratory values

We also found several pleiotropic genes which were statistically significantly associated with plasma viral load, triglyceride levels, and/or absolute basophil count (Fig 7). These included *ABCB4, ATAT1, C11orf74, C4B, C6orf48, CD2AP, CDK5RAP3, CNBD2, F2RL1, GPATCH4, GPR22, KCNJ15, KCTD7, PARM1, PCDHB3, RPS28, TTI2, USP19*. Some of them were located on chromosome 6, surrounding the major histocompatibility complex (MHC) region, while the rest scattered across the human genome. Meanwhile, we did not observe correlations among plasma viral load, triglyceride levels, or absolute basophil count. The strongest correlation was observed between plasma viral load and triglyceride levels (r^2^ = 0.24), suggesting only weak correlation, and correlations for the other pairs of laboratory values were approximately 0. Overall, there were potential pleiotropic genes for plasma viral load, triglyceride levels, and/or absolute basophil count in HIV-positive adults.

## Discussion

### Novel design of the simulation framework

In this report, we described a novel simulation framework for TWAS, and evaluated TWAS gene prioritization performance for genes with various degrees of tissue specificity. Our simulation results validated conclusions from several previous eQTL or TWAS studies [13-15,21], and also generated new findings that warrant attention in future TWAS. First, TWAS methods tested in this study all had well-controlled type I error rates (≤ 5%) for genes with any degrees of tissue-specificity. Second, single-tissue TWAS tended to have higher false positive rates of tissues. The phenomenon became more obvious when genes had more correlated expression levels across tissues. For tissue-specific genes, false positive rates of tissues could be controlled (≤ 5%) by adopting a more stringent multiple testing correction approach. However, for ubiquitously expressed genes, false positive rates of tissues remained significant (∼77%) even after a stringent multiple testing adjustment. Third, TWAS gene prioritization power was improved by eQTLs that were identified by jointly analyzing transcriptomic data across tissues. Fourth, for tissue-specific genes, single-tissue and cross-tissue gene-level association approaches had similar power. For ubiquitously expressed and similarly expressed genes, cross-tissue association approaches had greater power.

We further tested our simulation designs for how they would translate to real-world data by evaluating trait heritability in our simulated datasets. We found no apparent effect of MAF distribution on trait heritability under TWAS models. Instead, trait heritability increased with 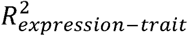. When 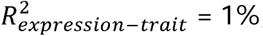, trait heritability was approximately 1% (s.e. = 0.059%). The estimated trait heritability was within a reasonable range and supported our simulation design.

### Associations in the clinical trials dataset

TSA-TWAS successfully replicated proof-of-concept gene-trait associations, including associations between *CETP* and HDL-c, and between *GPLD1* and plasma alkaline phosphatase levels. For total plasma bilirubin levels, our TSA-TWAS framework prioritized *UGT1A1* and genes near *UGT1A1. UGT1A1* encodes the hepatic protein that glucuronidates bilirubin [28], and has been known to affect bilirubin levels [24,25,27]. Other genes have been associated with total bilirubin levels in numerous studies [24-27]. These genes span 1Mbp at the 2q37.1 locus and are within the same topologically associating domain (TAD), which suggests that a regulatory mechanism may affect expression of the entire *KCNJ13*-*UGT1A*-*MROH2A* gene region.

TSA-TWAS has also identified several pleiotropic genes that linked plasma viral load, absolute basophil count, and/or triglyceride levels, which were otherwise independent from each other. Plasma viral load is a strong predictor of clinical outcome and is highly variable among people living with HIV. Individuals vary in their ability in suppressing viral loads, in the absence of antiretroviral treatments. Moreover, people living with HIV experience dyslipidemia to different degrees. Grunfeld *et al*. [39] found that AIDS patients experienced different lipid changes from HIV-infected patients without AIDS. The discovery of pleiotropic genes suggests the complexity of HIV pathogenesis and provides a future direction for research on the complex inter-individual variability among people living with HIV.

### Limitations & future directions

Our simulations revealed high false positive rates of tissues for single-tissue TWAS. The high false positive rates seen with single-tissue TWAS may be due to limited sample sizes for eQTL discovery. GTEx analysis has shown that discovery of tissue-specific eQTLs is contingent on the sample sizes of tissues [10]. Unfortunately, many tissues still have limited sample sizes for the identification of tissue-specific eQTLs. Consequently, single-tissue TWAS may not have ample power to prioritize potential trait-related tissues. Adopting stricter multiple testing adjustment strategies for single-tissue TWAS is one practical approach to help reduce false positive rates in prioritized tissues, but this will sacrifice power.

The evaluation of TWAS power and type I error rates estimated from this simulation study might be limited due to the small sample sizes (N = 2,000 for association analyses). We selected this sample size for simulation in order to make it comparable to the average sample size of the ACTG phase I-IV combined clinical traits interrogated in this study. TWAS gene prioritization power can be improve with greater sample, but also under influence of many other factors as shown in Veturi *et al*. [21] and this study. Thus, TWAS performance can differ from dataset to dataset when using different TWAS methods. It was difficult to take every factor into consideration in this work. We dedicated this study to explore tissue specificity’s impact on TWAS performance, and, for future TWAS studies, suggest customized simulation to better understand TWAS performance on specific datasets and diseases of interest.

## Conclusions

Gene-level association studies offer the opportunity to better understand the genetic architecture of complex human traits by leveraging regulatory information from both noncoding and coding regions of the genome. This may expedite translation of basic research discoveries to clinical applications. We provide a comprehensive simulation algorithm to fully investigate TWAS performance for diverse biological scenarios. Based on our simulation, we promote a TSA-TWAS analytic framework. TSA-TWAS framework on ACTG clinical trials data ascribed statistical significance to proof-of-concept gene-trait associations, and also found several novel associations and pleiotropic genes, suggesting the complexity of HIV-related traits that latest bioinformatics methods can reveal.

Additional work is needed to fully understand the tissue and genetic architecture underlying complex traits. The simulation algorithm and schema developed for this study is versatile enough to answer other questions regarding causal genes and tissues for complex traits. Overall, our work provides and tests a novel, flexible simulation framework and an TSA-TWAS analytic framework for future complex trait studies.

## Materials and Methods

### TWAS simulation design

The simulation study systematically evaluated how the tissue-specificity of eQTLs and gene expression levels influences TWAS gene prioritization performance. We assumed additive genetic effects of eQTLs on gene expression levels, and of gene expression levels on traits. The TWAS simulation scripts are available in R programming language at GitHub (https://github.com/BinglanLi/multi_tissue_twas_sim).

#### Genotype

We started by simulating genotypes for one gene in 1,500 individuals, which include eQTL and non-eQTL SNPs. Genotypes are denoted as *X*_*NxM*_ throughout this paper, where *N* denotes the total number of individuals and *M* denotes the total number of SNPs in a gene that include tissue-specific eQTLs, multi-tissue eQTLs and non-eQTL SNPs. These individuals were later stratified into an eQTL discovery dataset (*N*_*eQTL*_ = 500) and a TWAS testing dataset (*N*_*TWAS*_ = 1000), sample sizes comparable to those of current GTEx and ACTG datasets used in this analysis, respectively. Genotypes were simulated as biallelic SNPs and then converted into allele dosages as is done in most eQTL detection methods. MAF assigned to SNPs raged from 1% to 50% and were randomly drawn from a uniform distribution, *U*(0.01,0.5). Parameter settings of eQTLs in this simulation were drawn from observations in different eQTL databases (S1-3 Figs).

#### Gene expression level

We simulated one gene’s standardized expression levels at a time such that it was expressed in a fixed number of tissues. Let *P* denote the number of tissues where the gene is expressed, *P* = 1, 2, 5, or 10. If a gene is only expressed in a single tissue (*P* = 1), then, only single-tissue eQTLs were simulated for this given gene and no multi-tissue eQTLs were present.

A previous study showed that the number of eQTLs in a gene does not have as pronounced an effect on the TWAS power in comparison to other parameters [21]. Hence, we assumed that a given gene was regulated by the same total number of eQTLs in each of the *P* tissues, which is denoted by *M*_*eQTLs*_ (*M*_*eQTLs*_ = 30). eQTLs can be tissue-specific or have effect across multiple tissues. Here, we defined tissue-specific eQTLs as those that had effects in one and only one tissue. Multi-tissue eQTLs were defined as those who had effects in all *P* tissues in which the given gene is simulated to be expressed. We allowed multi-tissue eQTLs to have different effect sizes in different tissues. Assuming that a gene was expressed in *P* tissue(s) (say *P* = 5), then, this gene is regulated by both, tissue-specific eQTLs and multi-tissue eQTLs, in any of the *P* tissues. Let *M*_*ts-eQTLs*_ denote the number of tissue-specific eQTLs, and *M*_*mt-eQTLs*_ the number of multi-tissue eQTLs. A simulated gene had the same *M*_*ts-eQTLs*_ across *P* tissues, and the same *M*_*mt-eQTLs*_ across *P* tissues, such that *M*_*ts-eQTLs*_ and *M*_*mt-eQTLs*_ added up to *M*_*eQTLs*_ in each of the *P* tissues. Five different numbers of *M*_*mt-eQTLs*_ (0, 6, 12, 18, 24, corresponding *M*_*st-eQTLs*_ = 30, 24, 18, 12, 6) were evaluated, except when a gene was simulated to be expressed only in one gene, in which case *M*_*mt-eQTLs*_ always equaled 0.

Each gene was simulated under an additive genetic model per tissue. Let *E*_*N×P*_ denote the simulated gene expression levels for one gene, of *N* individuals, and across *P* tissues. For the given simulated gene, let *E*_*np*_ represent the simulated expression level of the *n*th individual in the *p*th tissue, which is an aggregate of tissue-specific eQTLs, multi-tissue eQTLs and non-eQTL effects in individual *n* for tissue *P*. The multivariate normal random effects model to simulate one gene’s expression levels is then expressed as follows:

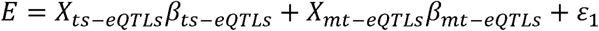

where *E* is the *N×P* matrix of standardized gene expression levels for a gene in *N* individuals across *P* tissues. *X*_*ts-eQTLs*_ is the *N×M*_*ts-eQTLs*_ matrix of standardized tissue-specific eQTL genotypes. Similarly,*X*_*mt-eQTLs*_ is the *N×M*_*ts mt-eQTLs*_ matrix of standardized multi-tissue eQTL genotypes. *β*_*ts-eQTLs*_ is a *M*_*ts-eQTLs*_ ×*P* matrix of tissue-specific eQTL effects. *β*_*ts-eQTLs,ip*_ represents the *i*th tissue-specific eQTL in the *p*th tissue, which could be a different eQTL across *P* tissues. Each value in the *β*_*ts-eQTLs*_ is independent of the others. *β*_*mt-eQTLs*_ is a *M*_*mt-eQTLs*_ ×*P* matrix of multi-tissue eQTL effects wherein *β*_*mt-eQTLs,jp*_ represents the *j*th multi-tissue eQTL in the *p*th tissue. In contrast to tissue-specific eQTLs, *β*_*mt-eQTLs,j*._ denotes the same *j*th multi-tissue eQTL in all *P* tissues, and is allowed to have similar or dissimilar effect sizes across *P* tissues (explained later in this section). 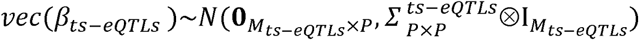 *where* 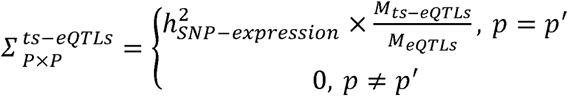. The constant, 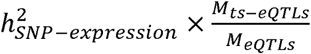, represents the proportion of variation in gene expression that can be explained by tissue-specific eQTLs. 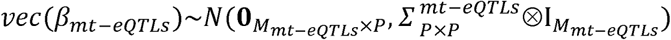 *where* 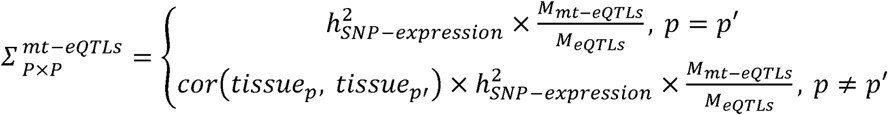. The constant, 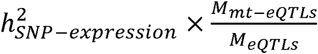, represents the proportion of gene expression variation that can be explained by multi-tissue eQTLs. *cor*(*tissue*_*p*_, *tissue*_*p′*_) represents the extent of similarity between *β*_*mt-eQTLs*,.*p*_ and *β*_*mt-eQTLs*,.*p′*_, i.e. the Pearson Correlation Coefficient between multi-tissue eQTL effect sizes in the *p*th and *p’*th tissues, respectively. The simulation algorithm allows multi-tissue eQTLs to have five different levels of *cor*(*tissue*_*i*_, *tissue*_*j*_) (0, 0.2, 0.4, 0.6, and 0.8). *ε*_1_ is the *N*×*P* matrix of residual errors that represent non-eQTL effects on a gene’s expression level and 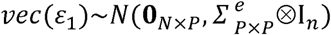 where 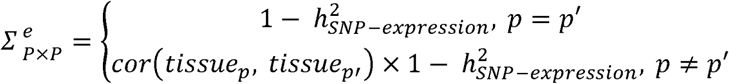. The constant, 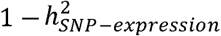, represents the proportion of gene expression variation that can be explained by factors other than eQTLs that can also regulate a gene’s final transcription isoforms and levels. We designed the error term to have such a covariance structure that the final aggregate expression levels of the given gene in *p*th tissue (*E*_.*p*_) was correlated with that in the *p’*th tissue (*E*_.*p′*_) due to multi-tissue eQTLs as well as other biological factors. These other biological factors (such as alternative splicing events, post-transcriptional modifications and regulation of mRNA degradation) can either be shared or different across tissues. We adopted a simple assumption that the more similar a gene’s expression levels are across tissues, the more likely multi-tissue eQTLs (and non-eQTL biological factors) will share effect sizes across tissues. Thus, correlation of gene expression across tissues (for example, correlation between *E*_.*p*_ and *E*_.*p′*_) is expected to be similar to, if not the same as, the correlation of multi-tissue eQTL effect sizes (for example, correlation between *β*_*mt-eQTLs*,.*p*_ and *β*_*mt-eQTLs*,.*p′*_) as well as the correlation between non-eQTL biological factors. All three random effect terms, i.e. *β*_*ts-eQTLs*_, *β*_*mt-eQTLs*_, and *ε*_1_ were simulated using the *rmvnorm* function from the R package, *mvtnorm*. We evaluated the extent of bias between assumed combination of simulation parameters and those estimated from the empirical distribution of simulated *E*_*N×P*_, which met the expectation (S16 Fig).

In the special case where a gene was simulated to be expressed only in a single tissue, the model was equivalent to a univariate normal distribution with mean 0 and variance equal to the expression heritability of that gene.

Tissue specificity of genes was characterized by the number of tissues in which genes are expressed as well as the similarity of gene expression levels across tissues. Tissue specificity of eQTLs was characterized by the proportion of multi-tissue eQTLs in a gene, the number of tissues where multi-tissue eQTLs were effective, and the similarity of eQTL effect sizes across tissues.

#### Phenotype

We assumed one and only one causal tissue for a phenotype and simulated phenotype datasets for the TWAS testing dataset (*N* = 1,000). This design was adopted from the simulation work of Dr. Yiming Hu *et al*. in the paper that described UTMOST [15]. Let *E*_*eQTLs*_ denote the standardized genetically regulated expression component in the causal tissue. The model to simulate traits from gene expression levels can be expressed as *Y* = *E*_*eQTLs*_ *b*_1_ + *ε*_2_, where *Y* is a 1000 × 1 vector of standardized responses for the 1,000 individuals in the TWAS testing dataset, *b*_1_ is the *M*_*eQTLs*_ ×1 vector of gene expression effect drawn from a normal distribution with mean zero and variance 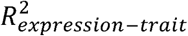, and *ε*_2_ is the vector of normally-distributed errors with mean zero and variance 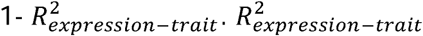 was assigned values in 0.001%, 0.05%, 0,5% and 1%, to represent different strengths of gene expression level-trait relations. To evaluate type I error rates, 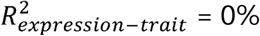 corresponded to the null model where gene and trait were unrelated.

#### eQTL detection

We adopted two types of eQTL detection methods, 1) elastic net (implemented in PrediXcan [5]) and 2) group LASSO (implemented in UTMOST [15]). For ease of parallel computation, these two algorithms were adapted and integrated into the TWAS simulation tool scripts. eQTLs detected in a single tissue context (elastic net) and those detected in an integrative tissue context (group LASSO) were then used to impute GReX, and for gene-trait association analyses.

#### Imputation of GreX

Expression level of a gene can be imputed using a linear model as *E* = *Xβ*, where *E* is the N×1 vector of imputed gene expression levels of the gene, *X* is the *N*×*M* matrix of genotypes, and *β* is the *M*×1 vector of eQTLs’ estimated regulatory effects on the gene, and can be obtained by either elastic net or group LASSO.

#### Association analysis

Single-tissue gene-trait associations were then estimated using SLR model, i.e., *lm* function in R. Cross-tissue gene-trait association analyses were also conducted in R but using PC Regression (implemented in MulTiXcan [16]) and GBJ test (implemented in UTMOST [15]).

#### Measures of TWAS performance

Each combination of simulation parameters was repeated 100 times independently to assess power and type I error rates at *α* = 0.05. Estimation of TWAS power was calculated as the percentage that a simulated causal gene was successfully identified as statistically significant in the causal tissue in the hundred simulations. Estimation of TWAS type I error rates were calculated as the percentage that a gene was falsely identified as statistically significant when there was no gene-trait signal simulated in the hundred simulations. We assumed that a gene is related to a trait in a single tissue, which is often the case for non-pleiotropic genes. In the simulation, we knew the causal tissues for the simulated traits. We calculated the false positive rates of tissues by counting the proportion of statistically significant results that were in non-causal tissues.

The entire process was repeated 20 times for each combination of simulation parameters to avoid sampling variability and to determine distributions of power, type I error rates, and false positive rates of tissues. We further evaluated the statistical significance of the differences in power and type I error rate between every pairs of TWAS methods using Wilcoxon Signed-rank test.

### Evaluation of simulated genetic scenarios

Trait heritability assessment validated and supported our design of simulation parameters. We designed a Monte Carlo simulation approach to randomly generate eQTL-gene-trait relations using the aforementioned simulation tool. Each replication simulated one genotypic dataset and one subsequent GReX profile for a gene. We simulated 30 non-eQTL and 30 eQTL SNPs for 5,000 individuals in which MAF followed a uniform distribution of 1-50% and eQTLs explained 30% of gene expression variation. The GReX profile was then used to generate 50 different traits using different random seeds. Thus, each simulated genotypic dataset had 50 estimated trait heritability values available; we took the average of these as the point estimate of the trait *h*^2^ for each genotypic dataset. GCTA [40] was not appropriate for our simulation as it assumes genome-wide genotypic data. Instead, we used the R package, *regress*, to estimate trait heritability in the simulated datasets. The entire process was repeated 30 times to generate a distribution of estimated trait heritability for a given combination of simulation parameters.

To determine the influence of MAF on trait heritability, we designed different ranges of MAF distributions. MAF of SNPs followed a uniform distribution of 1-50% as in the primary TWAS performance evaluation, and also 1-20% and 1-5%. We also simulated traits where 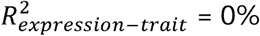 (negative control), 2%, or 5% (positive controls) to support the estimation of trait heritability when 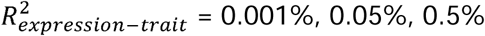, and 1%.

### AIDS Clinical Trials Group studies

The ACTG is the world’s largest HIV clinical trials network. It has conducted major clinical trials and translational biomedical research that have improved treatments and standards of care for people living with HIV in the United States and worldwide. In this study, we used data from four separate genotyping phases of specimens from ACTG studies in a combined dataset that comprises HIV treatment-naïve participants at least 18 years of age enrolled in randomized treatment trials [41-47]. Participants enrolled into ACTG protocols A5095, A5142, ACTG 384, A5202 or A5257. Informed consent for genetic research was obtained under ACTG protocol A5128. Clinical trial designs and outcomes, and results of a genome-wide pleiotropic study for baseline laboratory values have been described elsewhere[24,25].

### Genotypic data and quality control

A total of 4,411 individuals were genotyped in four phases. Phase I (samples from study A5095) was genotyped using Illumina 650Y array; Phase II (studies ACTG384 and A5142) and III (study A5202) were genotyped using Illumina 1M duo array; Phase IV (study 5257) was genotyped using Illumina HumanCoreExome BeadChip. Preparation of genotypic data included pre-imputation quality control (QC), imputation, and post-imputation QC. Pre- and post-imputation QC followed the same guidelines [48] and used PLINK1.90 [49] and R programming language. Imputation was performed on the combined ACTG phase I-IV genotype dataset after pre-imputation QC, which used IMPUTE2 [50] with 1000 Genomes Phase 1 v3 [51] as the reference panel. Combined ACTG phase I-IV imputed data comprised 27,438,241 variants. The following procedures/parameters were used in the post-imputation QC by PLINK1.90: sample inclusion in the ACTG genotyping phase I-IV phenotype collection, biallelic SNP check, imputation score (> 0.7), concordance of genetic and self-reported sex, genotype call rate (> 99%), sample call rate (> 98%), MAF (> 5%), and relatedness check (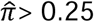; one individual was dropped from each related pair). Subsequent principal component analysis (EIGENSOFT [52]) projected remaining individuals onto the 1000 Genomes Project Phase 3 sample space to examine for population stratification. Based on percent of variance explained, the first three principal components estimated by SmartPCA in EIGENSOFT were used as covariates to adjust for population structure in the subsequent analyses. The final QC’ed ACTG phase I-IV combined imputed data comprised 2,185,490 genotyped and imputed biallelic SNPs for 4,360 individuals.

### Phenotypic data and QC

Data for 37 baseline (i.e., pre-treatment) laboratory measures were available from 5,185 HIV treatment-naive individuals in the ACTG genotyping phase I-IV datasets. We assembled these laboratory traits using a MySQL database and applied QC using R. We retained only individuals with available genotype data, and traits that were normally distributed and met the criterion of phenotype missing rate < 80%. Frequency distributions of traits were inspected using *hist_plot*.*R* that facilitates manual inspection of continuous traits by providing fast, high-throughput visualization along with necessary summary statistics of each visualized traits[53]. *hist_plot*.*R* is part of the CLARITE [53], which is available at https://github.com/HallLab/clarite. We also cross-referenced the retained traits to other published work that analyzed the same traits using these clinical trials datasets [24,25]. Non-fasting serum lipid measures were retained based on data from several studies [54-56]. The final combined dataset for ACTG genotyping phases I-IV comprised 37 baseline laboratory traits (Table 2).

### Description of a general TSA-TWAS analytic framework

The TSA-TWAS analytic framework has the following general steps.

1. Impute the GReX for the gene based on the input eQTL database(s) and the genotypic dataset.
2. Determine whether the gene is predicted to be expressed in only one tissue or in multiple tissues.
3. If the gene is predicted to be expressed in only one tissue, perform single-tissue TWAS using simple linear or logistic regression depending on the trait.
4. If the gene is predicted to be expressed in multiple tissues, perform cross-tissue TWAS using the GBJ test.
5. Repeat step 2-4 for the next gene.
6. (Optional) If there is more than one trait, repeat step 1-5 for the next trait.

### Imputation of GReX for genes

We used GTEx v8 MASHR-based eQTLs models [57] to impute gene expression levels in a tissue-specific manner. MASHR-based eQTLs models selected variants that have biological evidence of a potential causal role in gene expression, and estimated these variants’ effect sizes on gene expression levels in 49 tissues, using GTEx v8 as the reference dataset (available at http://predictdb.org/). The GTEx v8 MASHR-based eQTLs models were downloaded from their website on October 31, 2019. The QC’ed ACTG phase I-IV combined imputed data was used to impute the individual-level GReX in 49 human tissues.

### Statistical analysis for Gene-level associations

We tested for single-tissue gene-trait associations by performing association tests on imputed GReX and ACTG baseline lab traits using PLATO [58,59] in 49 tissues, separately. All baseline laboratory traits were continuous and thus were modeled by linear regression with covariates. Covariates included age, sex, and the first three principal components calculated by EIGENSOFT to adjust for sampling biases and underlying population structure. For cross-tissue association analyses, we adapted the UTMOST script in R programming language and performed the GBJ test for the individual-level ACTG data. The lowest p-value that can be generated by GBJ test in R is approximately 1×10^−15^. No obvious inflation was observed in the TSA-TWAS framework. ACTG phenome-wide TWAS results were visualized using PhenoGram [60], a web-based, versatile data visualization tool to create chromosomal ideograms with customized annotations, available at http://visualization.ritchielab.org/phenograms/plot. Supplementary manhattan plot was created by *hudson*, a R package available at https://github.com/anastasia-lucas/hudson.

### Statistical correction

Two strategies to correct for multiple testing were implemented in the ACTG analysis, method-wise and family-wise Bonferroni significance thresholds. The method-wise approach ascribes significance to statistical tests by controlling for the number of tests conducted in one type of method. For single-tissue gene-trait associations, the method-wise Bonferroni significance threshold was corrected for the number of genes (n = 483) and traits (n = 37), which resulted in 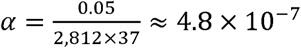. For cross-tissue gene-trait associations, the method-wise Bonferroni significance threshold corrected for the number of genes and traits, which gave 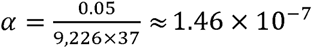. The family-wise approach assigns significance to tests by accounting for all tests performed in this study to control for FWER. Hence, single-tissue and cross-tissue association tests shared the same family-wise Bonferroni significance threshold, 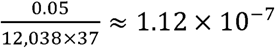. The significance threshold for interpreting results, by default, referred to the family-wise threshold. All results reported are exact p-values and thus, can be easily compared to either multiple testing threshold.

## Acknowledgements

The authors are grateful to the many persons living with HIV who volunteered for ACTG protocols A5095, A5142, ACTG 384, A5202, and A5257. In addition, they acknowledge the contributions of study teams and site staff for these protocols. We thank Paul J. McLaren, PhD (Public Health Agency of Canada, Winnipeg, Canada) for prior involvement and collaborations that used these genome-wide genotype data. Study drugs were provided by DuPont Pharmaceutical Company, Bristol-Myers Squibb, Inc., Agouron Pharmaceuticals, Inc., GlaxoWellcome, Inc., Merck and Co., Inc. Boehringer-Ingelheim Pharmaceuticals, Inc., Gilead Sciences, Inc., GlaxoSmithKline, Inc., Abbott Laboratories, Inc., Tibotec Therapeutics. The clinical trials were ACTG 384 (ClinicalTrials.gov: NCT00000919), A5095 (NCT00013520), A5142 (NCT00050895), A5202 (NCT00118898), and A5257 (NCT00811954). We thank Dr. Yiming Hu and Zhaolong Yu from Yale University and Alvaro Barbeira and Dr. Hae Kyung Im from the University of Chicago for technical support. We thank Dr. Shefali S. Verma and Anastasia Lucas from Dr. Marylyn Ritchie’s lab at the University of Pennsylvania for suggestions and assistance in data visualization.

This project was supported by the National Institute of Allergy and Infectious Diseases (NIAID award number U01AI068636), the National Institute of Mental Health, and the National Institute of Dental and Craniofacial Research.

Grant support included TR000124 (to E.S.D.); AI077505, TR000445, AI069439 (to D.W.H.); and the National Institute of Allergy and Infectious Disease (NIAID award AI077505 and AI116794 (to M.D.R.).

Clinical research sites that participated in ACTG protocols ACTG 384, A5095, A5142, A5202 or A5257, and collected DNA under protocol A5128 were supported by the following grants from the National Institutes of Health (NIH): A1069412, A1069423, A1069424, A1069503, AI025859, AI025868, AI027658, AI027661, AI027666, AI027675, AI032782, AI034853, AI038858, AI045008, AI046370, AI046376, AI050409, AI050410, AI050410, AI058740, AI060354, AI068636, AI069412, AI069415, AI069418, AI069419, AI069423, AI069424, AI069428, AI069432, AI069432, AI069434, AI069439, AI069447, AI069450, AI069452, AI069465, AI069467, AI069470, AI069471, AI069472, AI069474, AI069477, AI069481, AI069484, AI069494, AI069495, AI069496, AI069501, AI069501, AI069502, AI069503, AI069511, AI069513, AI069532, AI069534, AI069556, AI072626, AI073961, RR000046, RR000425, RR023561, RR024156, RR024160, RR024996, RR025008, RR025747, RR025777, RR025780, TR000004, TR000058, TR000124, TR000170, TR000439, TR000445, TR000457, TR001079, TR001082, TR001111, and TR024160.

## Reference

1. Visscher PM, Wray NR, Zhang Q, Sklar P, McCarthy MI, Brown MA, et al. 10 Years of GWAS Discovery: Biology, Function, and Translation. The American Journal of Human Genetics. ElsevierCompany; 2017 Jul 6;101(1):5–22.

2. Lappalainen T. Functional genomics bridges the gap between quantitative genetics and molecular biology. Genome Research. Cold Spring Harbor Lab; 2015 Oct;25(10):1427–31.

3. MacArthur J, Bowler E, Cerezo M, Gil L, Hall P, Hastings E, et al. The new NHGRI-EBI Catalog of published genome-wide association studies (GWAS Catalog). Nucleic Acids Research. 2017 Jan 4;45(D1):D896–D901.

4. Maurano MT, Humbert R, Rynes E, Thurman RE, Haugen E, Wang H, et al. Systematic localization of common disease-associated variation in regulatory DNA. Science. American Association for the Advancement of Science; 2012 Sep 7;337(6099):1190–5.

5. Gamazon ER, Wheeler HE, Shah KP, Mozaffari SV, Aquino-Michaels K, Carroll RJ, et al. A gene-based association method for mapping traits using reference transcriptome data. Nat Genet. Nature Publishing Group; 2015 Aug 10;47(9):1091–8.

6. Gusev A, Ko A, Shi H, Bhatia G, Chung W, Penninx BWJH, et al. Integrative approaches for large-scale transcriptome-wide association studies. Nat Genet. Nature Publishing Group; 2016 Mar;48(3):245–52.

7. Thériault S, Gaudreault N, Lamontagne M, Rosa M, Boulanger M-C, Messika-Zeitoun D, et al. A transcriptome-wide association study identifies PALMD as a susceptibility gene for calcific aortic valve stenosis. Nature Communications. Nature Publishing Group; 2018 Mar 7;9(1):988.

8. Wu L, Shi W, Long J, Guo X, Michailidou K, Beesley J, et al. A transcriptome-wide association study of 229,000 women identifies new candidate susceptibility genes for breast cancer. Nat Genet. Nature Publishing Group; 2018 Jun 18;50(7):968–78.

9. Mancuso N, Shi H, Goddard P, Kichaev G, Gusev A, Pasaniuc B. Integrating Gene Expression with Summary Association Statistics to Identify Genes Associated with 30 Complex Traits. American journal of human genetics. Elsevier; 2017 Mar 2;100(3):473–87.

10. Battle A, Brown CD, Engelhardt BE, Montgomery SB. Genetic effects on gene expression across human tissues. Nature. Nature Publishing Group; 2017 Oct 12;550(7675):204–13.

11. Võsa U, Claringbould A, Westra H-J, Bonder MJ, Deelen P, Zeng B, et al. Unraveling the polygenic architecture of complex traits using blood eQTL meta-analysis. 2018.

12. Li B, Veturi Y, Bradford Y, Verma SS, Verma A, Lucas AM, et al. Influence of tissue context on gene prioritization for predicted transcriptome-wide association studies. Pac Symp Biocomput. 2019;24:296–307.

13. Flutre T, Wen X, Pritchard J, Stephens M. A statistical framework for joint eQTL analysis in multiple tissues. Gibson G, editor. PLoS Genet. Public Library of Science; 2013 May;9(5):e1003486.

14. Sul JH, Han B, Ye C, Choi T, Eskin E. Effectively identifying eQTLs from multiple tissues by combining mixed model and meta-analytic approaches. Schork NJ, editor. PLoS Genet. Public Library of Science; 2013 Jun;9(6):e1003491.

15. Hu Y, Li M, Lu Q, Weng H, Wang J, Zekavat SM, et al. A statistical framework for cross-tissue transcriptome-wide association analysis. Nat Genet. Nature Publishing Group; 2019 Mar;51(3):568–76.

16. Barbeira AN, Pividori M, Zheng J, Wheeler HE, Nicolae DL, Im HK. Integrating predicted transcriptome from multiple tissues improves association detection. Plagnol V, editor. PLoS Genet. 2019 Jan;15(1):e1007889.

17. Liu X, Finucane HK, Gusev A, Bhatia G, Gazal S, O’Connor L, et al. Functional Architectures of Local and Distal Regulation of Gene Expression in Multiple Human Tissues. American journal of human genetics. 2017 Apr 6;100(4):605–16.

18. Wainberg M, Sinnott-Armstrong N, Mancuso N, Barbeira AN, Knowles DA, Golan D, et al. Opportunities and challenges for transcriptome-wide association studies. Nat Genet. Nature Publishing Group; 2019 Apr;51(4):592–9.

19. Fagerberg L, Hallström BM, Oksvold P, Kampf C, Djureinovic D, Odeberg J, et al. Analysis of the human tissue-specific expression by genome-wide integration of transcriptomics and antibody-based proteomics. Mol Cell Proteomics. American Society for Biochemistry and Molecular Biology; 2014 Feb;13(2):397–406.

20. Kryuchkova-Mostacci N, Robinson-Rechavi M. Tissue-Specificity of Gene Expression Diverges Slowly between Orthologs, and Rapidly between Paralogs. Ouzounis CA, editor. PLoS Comput Biol. Public Library of Science; 2016 Dec;12(12):e1005274.

21. Veturi Y, Ritchie MD. How powerful are summary-based methods for identifying expression-trait associations under different genetic architectures? Pac Symp Biocomput. 2018;23:228–39.

22. Wheeler HE, Shah KP, Brenner J, Garcia T, Aquino-Michaels K, GTEx Consortium, et al. Survey of the Heritability and Sparse Architecture of Gene Expression Traits across Human Tissues. Montgomery SB, editor. PLoS Genet. 2016 Nov 11;12(11):e1006423–3.

23. Ongen H, Brown AA, Delaneau O, Panousis NI, Nica AC, GTEx Consortium, et al. Estimating the causal tissues for complex traits and diseases. Nat Genet. 2017 Dec;49(12):1676–83.

24. Moore CB, Verma A, Pendergrass S, Verma SS, Johnson DH, Daar ES, et al. Phenome-wide Association Study Relating Pretreatment Laboratory Parameters With Human Genetic Variants in AIDS Clinical Trials Group Protocols. Open Forum Infect Dis. 2015 Jan;2(1):ofu113.

25. Verma A, Bradford Y, Verma SS, Pendergrass SA, Daar ES, Venuto C, et al. Multiphenotype association study of patients randomized to initiate antiretroviral regimens in AIDS Clinical Trials Group protocol A5202. Pharmacogenetics and Genomics. 2017 Mar;27(3):101–11.

26. Coltell O, Asensio EM, Sorlí JV, Barragán R, Fernández-Carrión R, Portolés O, et al. Genome-Wide Association Study (GWAS) on Bilirubin Concentrations in Subjects with Metabolic Syndrome: Sex-Specific GWAS Analysis and Gene-Diet Interactions in a Mediterranean Population. Nutrients. Multidisciplinary Digital Publishing Institute; 2019 Jan 4;11(1):90.

27. Dai X, Wu C, He Y, Gui L, Zhou L, Guo H, et al. A genome-wide association study for serum bilirubin levels and gene-environment interaction in a Chinese population. Genet Epidemiol. 2013 Apr;37(3):293–300.

28. Tukey RH, Strassburg CP. Human UDP-glucuronosyltransferases: metabolism, expression, and disease. Annu Rev Pharmacol Toxicol. 2000;40:581–616.

29. Barter PJ, H Bryan Brewer J, Chapman MJ, Hennekens CH, Rader DJ, Tall AR. Cholesteryl Ester Transfer Protein. Arterioscler Thromb Vasc Biol. Lippincott Williams & Wilkins; 2003 Feb 1;23(2):160–7.

30. Chambers JC, Zhang W, Sehmi J, Li X, Wass MN, Van der Harst P, et al. Genome-wide association study identifies loci influencing concentrations of liver enzymes in plasma. Nat Genet. 2011 Oct 16;43(11):1131–8.

31. Kanai M, Akiyama M, Takahashi A, Matoba N, Momozawa Y, Ikeda M, et al. Genetic analysis of quantitative traits in the Japanese population links cell types to complex human diseases. Nat Genet. Nature Publishing Group; 2018 Mar;50(3):390–400.

32. Le Clerc S, Coulonges C, Delaneau O, van Manen D, Herbeck JT, Limou S, et al. Screening low-frequency SNPS from genome-wide association study reveals a new risk allele for progression to AIDS. J Acquir Immune Defic Syndr. 2011 Mar 1;56(3):279–84.

33. Rhee EP, Ho JE, Chen M-H, Shen D, Cheng S, Larson MG, et al. A genome-wide association study of the human metabolome in a community-based cohort. Cell Metab. 2013 Jul 2;18(1):130–43.

34. Lingwood CA, Branch DR. The role of glycosphingolipids in HIV/AIDS. Discov Med. Discov Med; 2011 Apr;11(59):303–13.

35. van Til NP, Heutinck KM, van der Rijt R, Paulusma CC, van Wijland M, Markusic DM, et al. Alteration of viral lipid composition by expression of the phospholipid floppase ABCB4 reduces HIV vector infectivity. Retrovirology. BioMed Central; 2008 Feb 1;5(1):14–9.

36. Wu B, Ouyang Z, Lyon CJ, Zhang W, Clift T, Bone CR, et al. Plasma Levels of Complement Factor I and C4b Peptides Are Associated with HIV Suppression. ACS Infect Dis. 2017 Dec 8;3(12):880–5.

37. Dunn SJ, Khan IH, Chan UA, Scearce RL, Melara CL, Paul AM, et al. Identification of cell surface targets for HIV-1 therapeutics using genetic screens. Virology. 2004 Apr 10;321(2):260–73.

38. Migueles SA, Sabbaghian MS, Shupert WL, Bettinotti MP, Marincola FM, Martino L, et al. HLA B*5701 is highly associated with restriction of virus replication in a subgroup of HIV-infected long term nonprogressors. PNAS. 2000 Mar 14;97(6):2709–14.

39. Grunfeld C, Pang M, Doerrler W, Shigenaga JK, Jensen P, Feingold KR. Lipids, lipoproteins, triglyceride clearance, and cytokines in human immunodeficiency virus infection and the acquired immunodeficiency syndrome. J Clin Endocrinol Metab. 1992 May;74(5):1045–52.

40. Yang J, Lee SH, Goddard ME, Visscher PM. GCTA: a tool for genome-wide complex trait analysis. American journal of human genetics. 2011 Jan 7;88(1):76–82.

41. Robbins GK, De Gruttola V, Shafer RW, Smeaton LM, Snyder SW, Pettinelli C, et al. Comparison of sequential three-drug regimens as initial therapy for HIV-1 infection. N Engl J Med. Massachusetts Medical Society; 2003 Dec 11;349(24):2293–303.

42. Gulick RM, Ribaudo HJ, Shikuma CM, Lustgarten S, Squires KE, Meyer WA, et al. Triple-nucleoside regimens versus efavirenz-containing regimens for the initial treatment of HIV-1 infection. N Engl J Med. Massachusetts Medical Society; 2004 Apr 29;350(18):1850–61.

43. Gulick RM, Ribaudo HJ, Shikuma CM, Lalama C, Schackman BR, Meyer WA, et al. Three-vs four-drug antiretroviral regimens for the initial treatment of HIV-1 infection: a randomized controlled trial. JAMA. 2006 Aug 16;296(7):769–81.

44. Riddler SA, Haubrich R, DiRienzo AG, Peeples L, Powderly WG, Klingman KL, et al. Class-sparing regimens for initial treatment of HIV-1 infection. N Engl J Med. Massachusetts Medical Society; 2008 May 15;358(20):2095–106.

45. Sax PE, Tierney C, Collier AC, Fischl MA, Mollan K, Peeples L, et al. Abacavir-lamivudine versus tenofovir-emtricitabine for initial HIV-1 therapy. N Engl J Med. Massachusetts Medical Society; 2009 Dec 3;361(23):2230–40.

46. Daar ES, Tierney C, Fischl MA, Sax PE, Mollan K, Budhathoki C, et al. Atazanavir Plus Ritonavir or Efavirenz as Part of a 3-Drug Regimen for Initial Treatment of HIV-1: A Randomized Trial. Ann Intern Med. American College of Physicians; 2011 Apr 5;154(7):445–56.

47. Lennox JL, Landovitz RJ, Ribaudo HJ, Ofotokun I, Na LH, Godfrey C, et al. A Phase III Comparative Study of the Efficacy and Tolerability of Three Non-Nucleoside Reverse Transcriptase Inhibitor-Sparing Antiretroviral Regimens for Treatment-Naïve HIV-1-Infected Volunteers: A Randomized, Controlled Trial. Ann Intern Med. NIH Public Access; 2014 Oct 7;161(7):461–71.

48. Turner S, Armstrong LL, Bradford Y, Carlson CS, Crawford DC, Crenshaw AT, et al. Quality control procedures for genome-wide association studies. Haines JL, Korf BR, Morton CC, Seidman CE, Seidman JG, Smith DR, editors. Curr Protoc Hum Genet. 2011 Jan;Chapter 1(1):Unit1.19–1.19.18.

49. Purcell S, Neale B, Todd-Brown K, Thomas L, Ferreira MAR, Bender D, et al. PLINK: a tool set for whole-genome association and population-based linkage analyses. The American Journal of Human Genetics. 2007 Sep;81(3):559–75.

50. Howie BN, Donnelly P, Marchini J. A Flexible and Accurate Genotype Imputation Method for the Next Generation of Genome-Wide Association Studies. Schork NJ, editor. PLoS Genet. Public Library of Science; 2009 Jun 19;5(6):e1000529.

51. 1000 Genomes Project Consortium, Abecasis GR, Altshuler D, Auton A, Brooks LD, Durbin RM, et al. A map of human genome variation from population-scale sequencing. Nature Publishing Group. 2010 Oct 28;467(7319):1061–73.

52. Price AL, Patterson NJ, Plenge RM, Weinblatt ME, Shadick NA, Reich D. Principal components analysis corrects for stratification in genome-wide association studies. Nat Genet. Nature Publishing Group; 2006 Aug;38(8):904–9.

53. Lucas AM, Palmiero NE, McGuigan J, Passero K, Zhou J, Orie D, et al. CLARITE Facilitates the Quality Control and Analysis Process for EWAS of Metabolic-Related Traits. Front Genet. Frontiers; 2019 Dec 18;10:1164.

54. Langsted A, Nordestgaard BG. Nonfasting versus fasting lipid profile for cardiovascular risk prediction. Pathology. 2019 Feb;51(2):131–41.

55. Nordestgaard BG. A Test in Context: Lipid Profile, Fasting Versus Nonfasting. J Am Coll Cardiol. 2017 Sep 26;70(13):1637–46.

56. Mora S, Chang CL, Moorthy MV, Sever PS. Association of Nonfasting vs Fasting Lipid Levels With Risk of Major Coronary Events in the Anglo-Scandinavian Cardiac Outcomes Trial-Lipid Lowering Arm. JAMA Intern Med. American Medical Association; 2019 May 28;179(7):898–905.

57. Barbeira AN, Bonazzola R, Gamazon ER, Liang Y, Park Y, Kim-Hellmuth S, et al. Exploiting the GTEx resources to decipher the mechanisms at GWAS loci. bioRxiv. Cold Spring Harbor Laboratory; 2020 May 23;42(D1):814350.

58. Hall MA, Wallace J, Lucas A, Kim D, Basile AO, Verma SS, et al. PLATO software provides analytic framework for investigating complexity beyond genome-wide association studies. Nature Communications. Nature Publishing Group; 2017 Oct 27;8(1):1167.

59. Grady BJ, Torstenson E, Dudek SM, Giles J, Sexton D, Ritchie MD. Finding unique filter sets in PLATO: a precursor to efficient interaction analysis in GWAS data. Pac Symp Biocomput. 2010;:315–26.

60. Wolfe D, Dudek S, Ritchie MD, Pendergrass SA. Visualizing genomic information across chromosomes with PhenoGram. BioData Min. BioMed Central; 2013 Oct 16;6(1):18–12.

