## Supplementary material for "Tissue specificity-aware TWAS (TSA-TWAS) framework identifies novel associations with metabolic, immunologic, and virologic traits in HIV-positive adults": S1 Fig

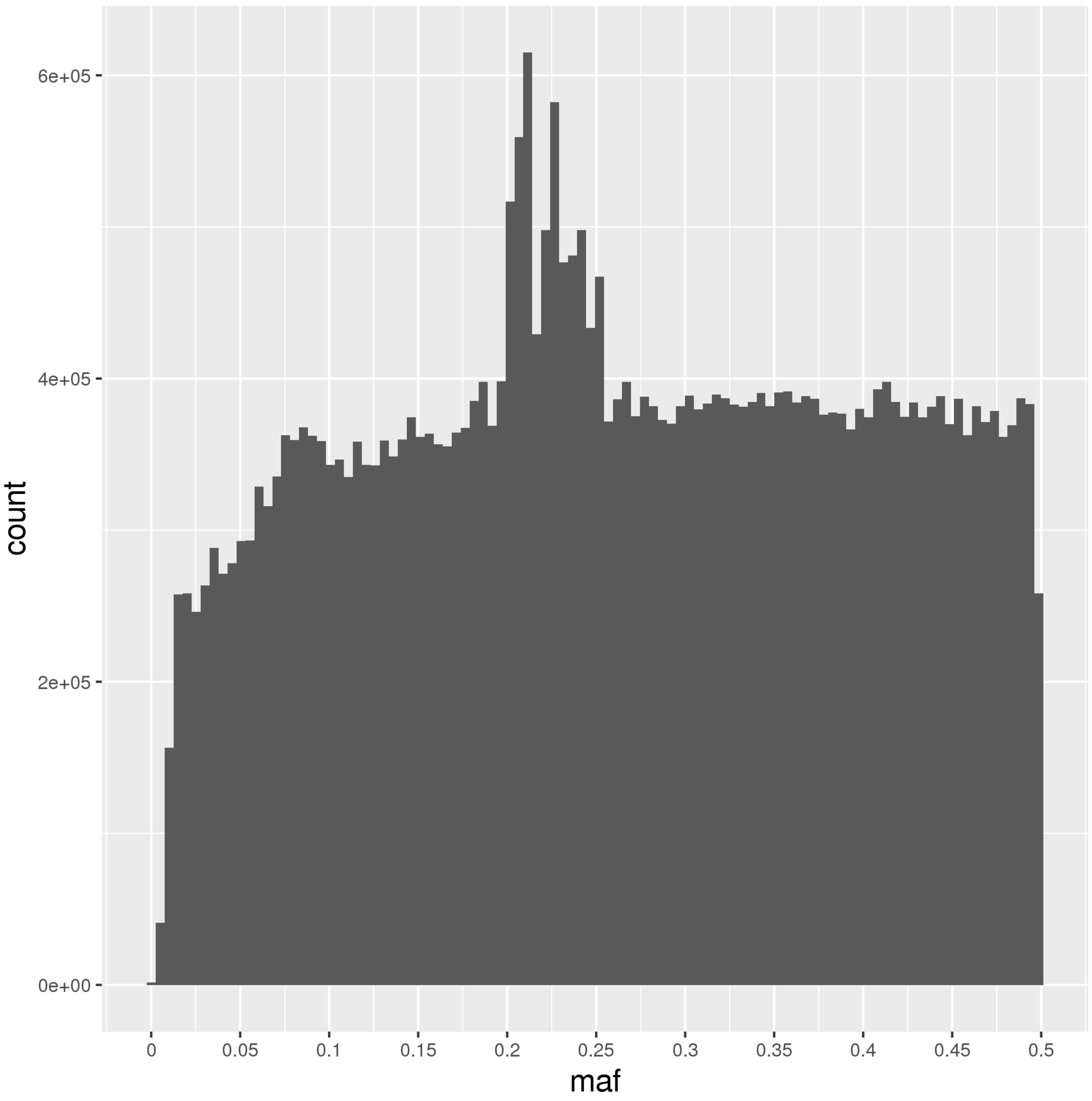

Figure S1. MAF distribution of GTEx v7 eQTLs. MAF of eQTLs closely resembled a uniform distribution, ranging between 1% to 50%, with a spike around 20-25%.
