## Supplementary material for "Tissue specificity-aware TWAS (TSA-TWAS) framework identifies novel associations with metabolic, immunologic, and virologic traits in HIV-positive adults": S2 Fig

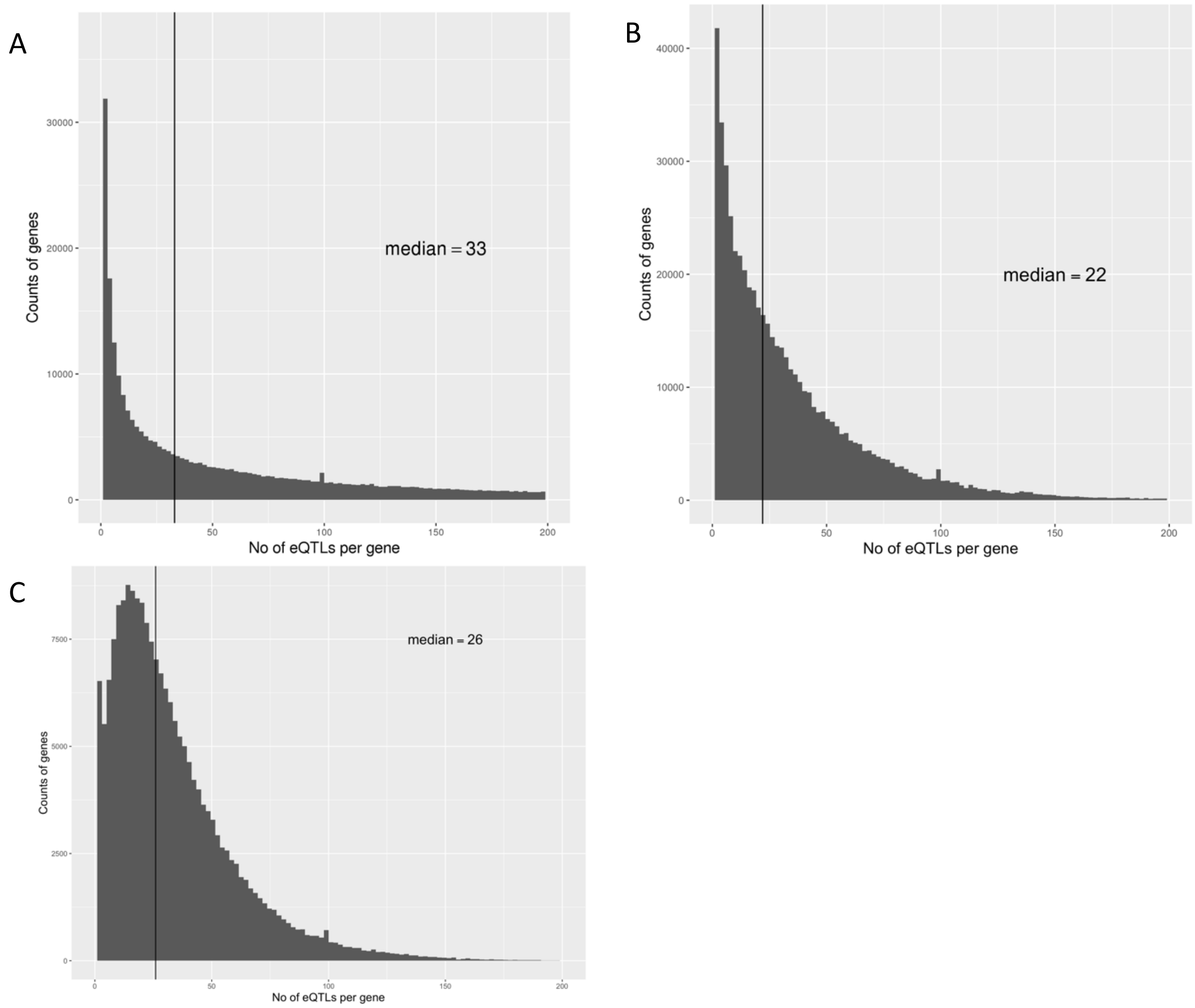


Figure S2. Distribution of number of eQTLs for a gene from **A)** GTEx v7, **B)** PredictDB eQTL datasets and **C)** UTMOST eQTL datasets.
