## Supplementary material for "Tissue specificity-aware TWAS (TSA-TWAS) framework identifies novel associations with metabolic, immunologic, and virologic traits in HIV-positive adults": S3 Fig

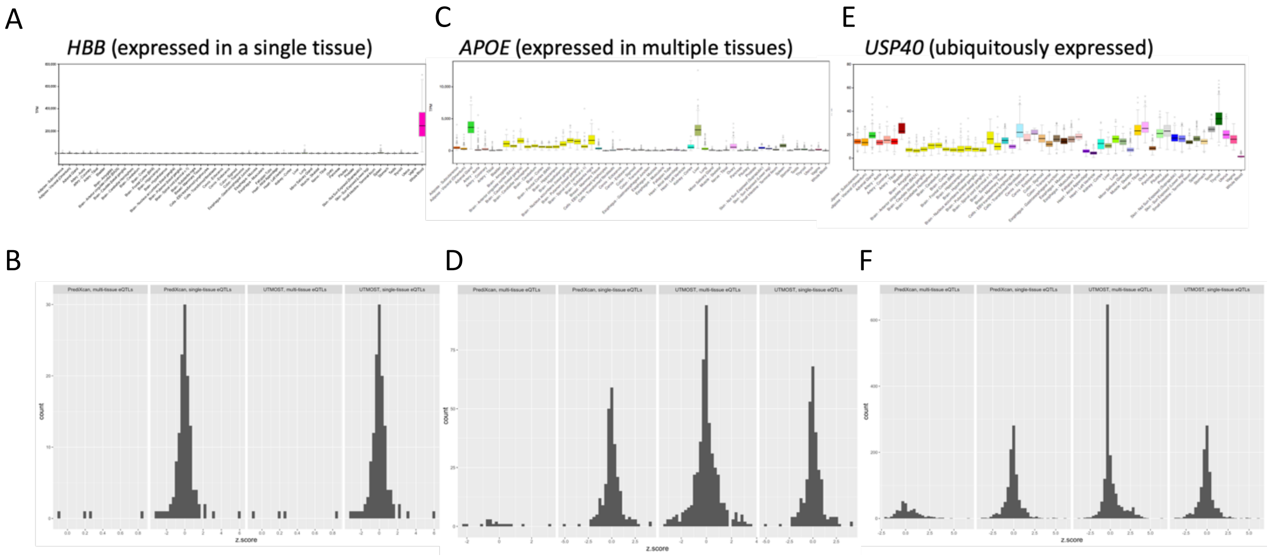


Figure S3. eQTL weight distribution for genes of different levels of tissue specificity. **A)** and **B)** For tissue-specific genes, like *HBB*, PrediXcan and UTMOST both identified eQTLs predominantly in a single tissue and eQTL weights followed a normal distribution. **C)** and **D)** For genes that are differentially expressed in multiple tissue, like *APOE*, PrediXcan and UTMOST both estimated normally distributed eQTLs weights. However, UTMOST was able to identify more eQTLs that are functioning across tissues. **E)** and **F)** For genes that are ubiquitously expressed in all tissues, like USP40, eQTL weights estimated from PrediXcan and UTMOST were both normally distributed. And again, UTMOST was able to identify more eQTLs that are effective in more than one tissues.
