## Supplementary material for "Tissue specificity-aware TWAS (TSA-TWAS) framework identifies novel associations with metabolic, immunologic, and virologic traits in HIV-positive adults": S4 Fig

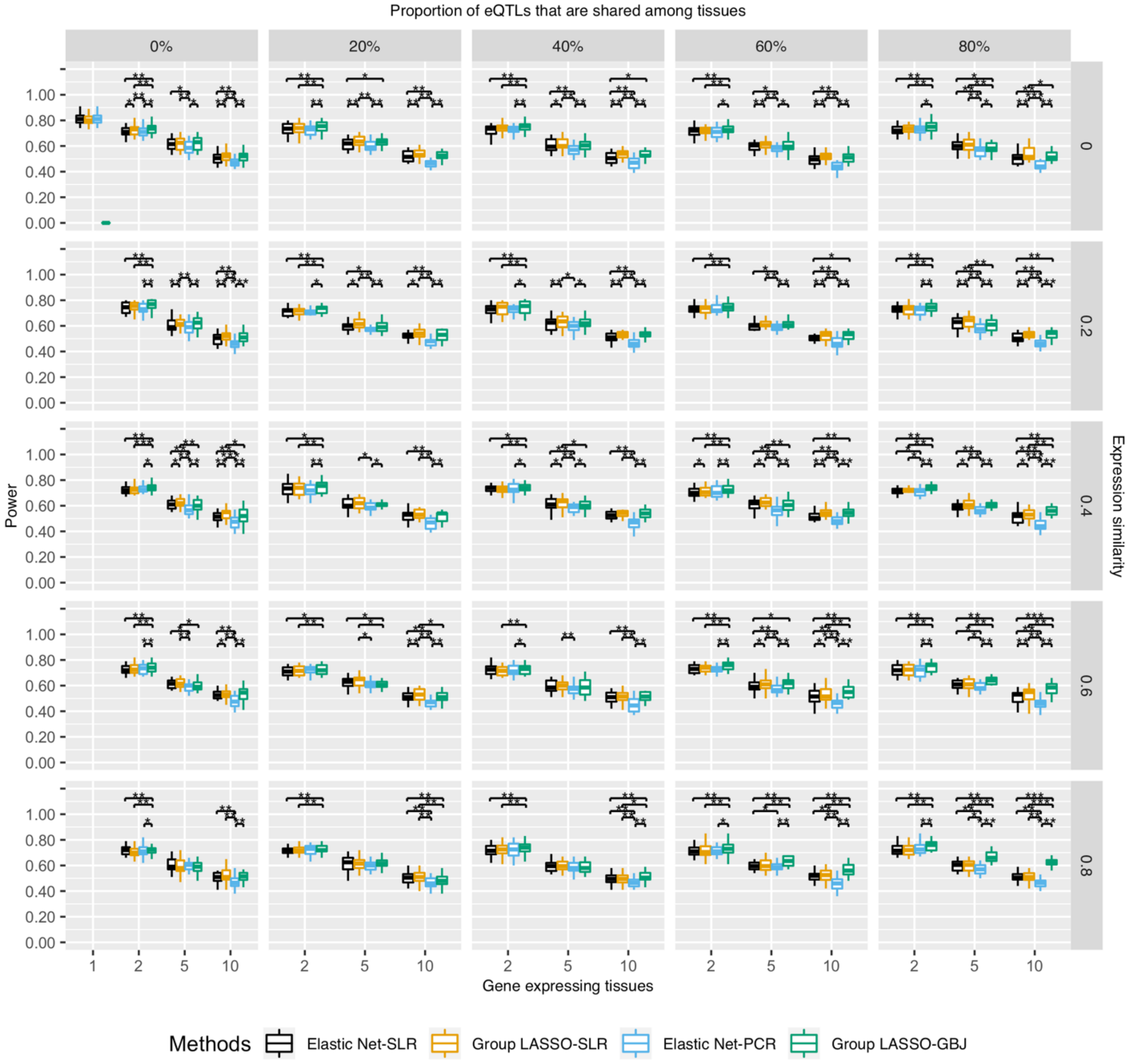


Figure S4. Statistical difference of gene prioritization power of different TWAS methods. Power was the proportion of successfully prioritized gene-trait associations in the causal tissue. X-axis is the number of gene-expressing tissues. Each column stands for the proportion of eQTLs that are shared among tissues for a gene. Each row is the similarity of gene expression profiles across tissues which is estimated by correlation. Moving from the top left to the bottom right is a gradient spectrum from tissue-specific genes to broadly expressed genes. The colors represent different TWAS methods and y-axis is the power. For tissue-specific genes at the top left, single-tissue TWAS (Elastic Net-SLR) and cross-tissue TWAS (Group LASSO-GBJ) had similar power. For broadly expressed genes at the bottom right, cross-tissue TWAS (Group LASSO-GBJ) had greater power. The difference in power among different TWAS methods were statistically evaluated (* p-value < 0.05, ** p-value < 0.01, *** p-value < 0.0001).
