## Supplementary figures and images for "Tissue specificity-aware TWAS (TSA-TWAS) framework identifies novel associations with metabolic, immunologic, and virologic traits in HIV-positive adults"

### S5 Fig

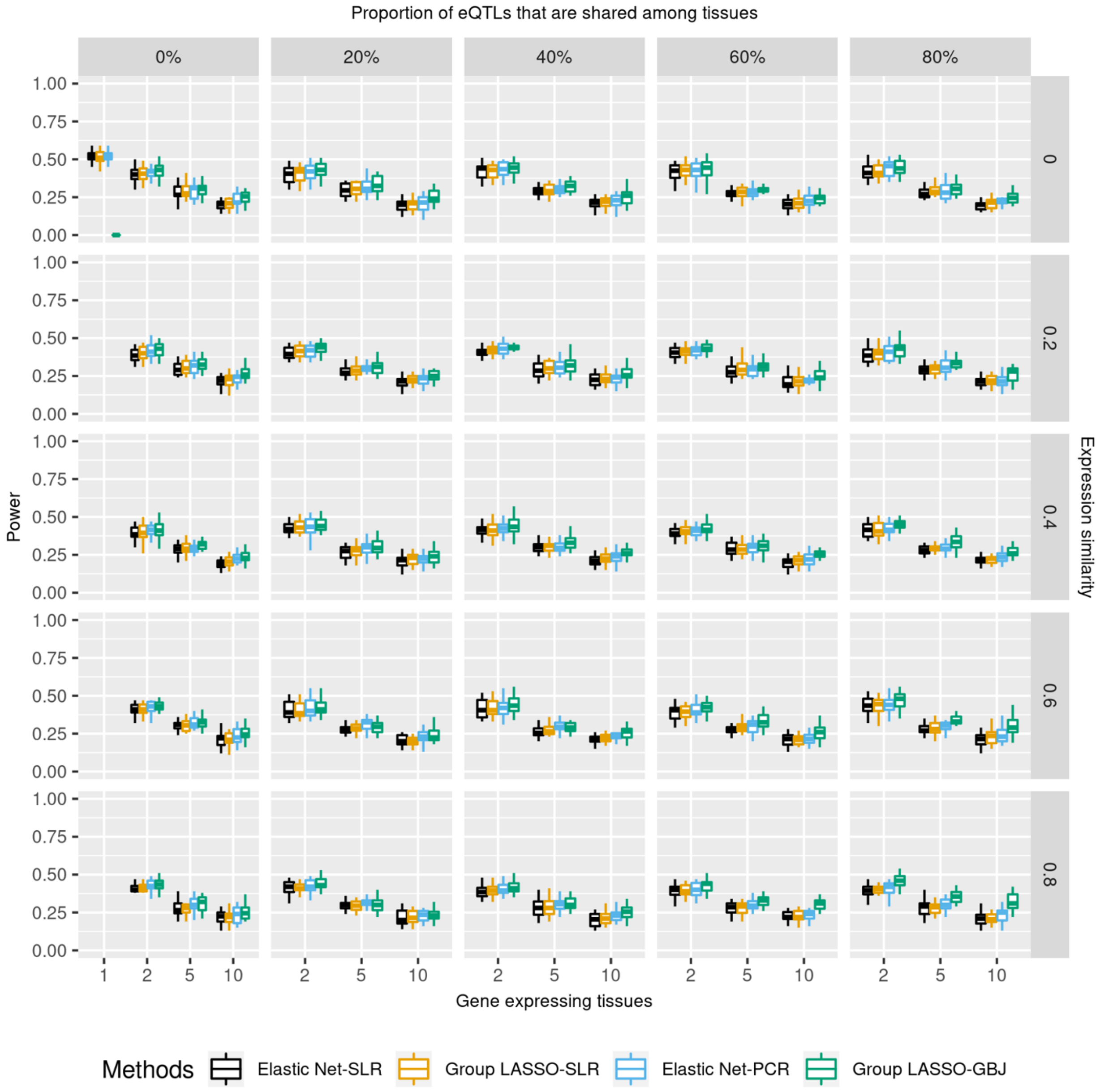


Figure S5. TWAS gene prioritization power when $R^2_{expression-trait}$ = 0.5%

### S6 Fig

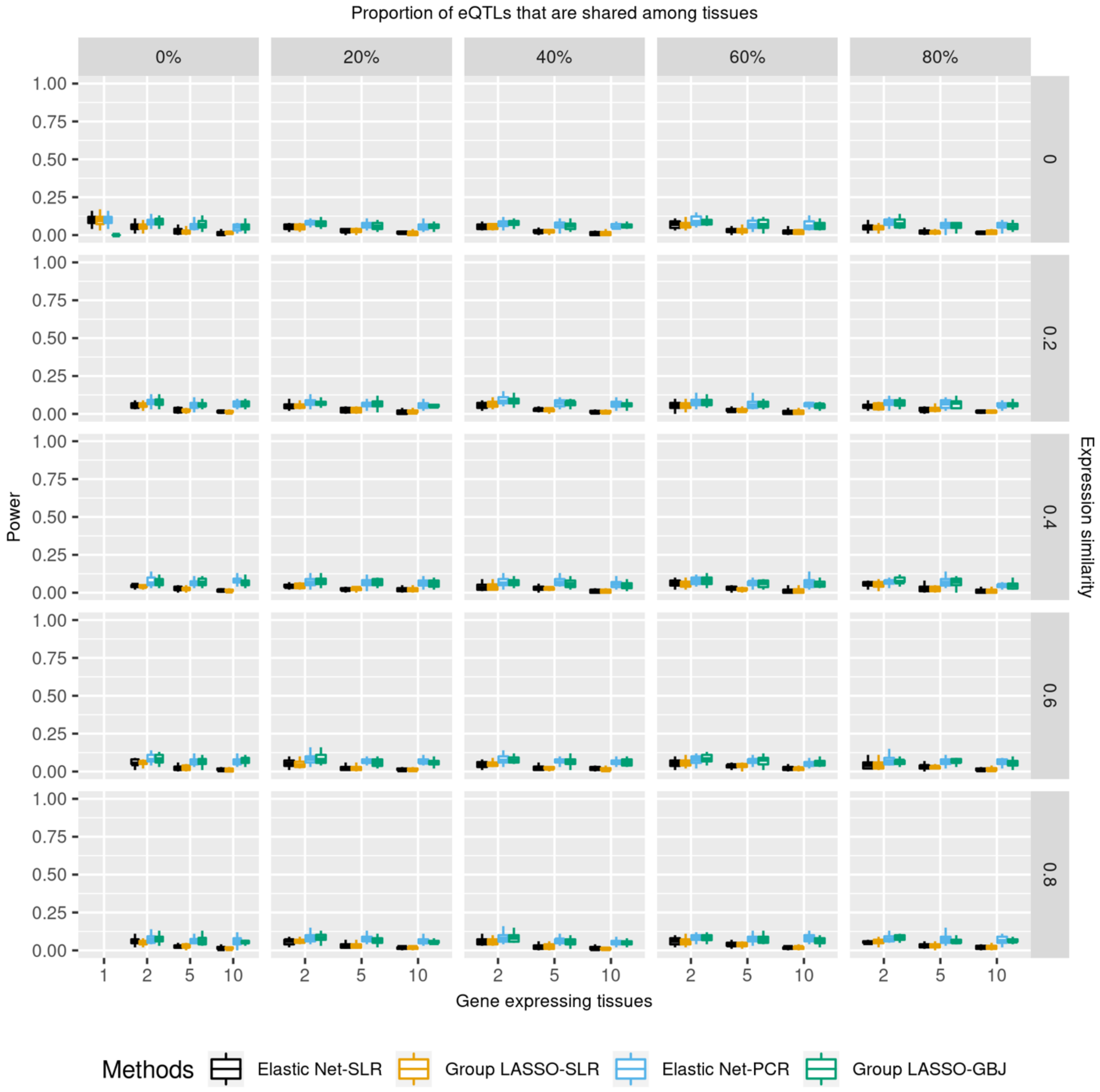


Figure S6. TWAS gene prioritization power when $R^2_{expression-trait}$ = 0.05%

### S7 Fig

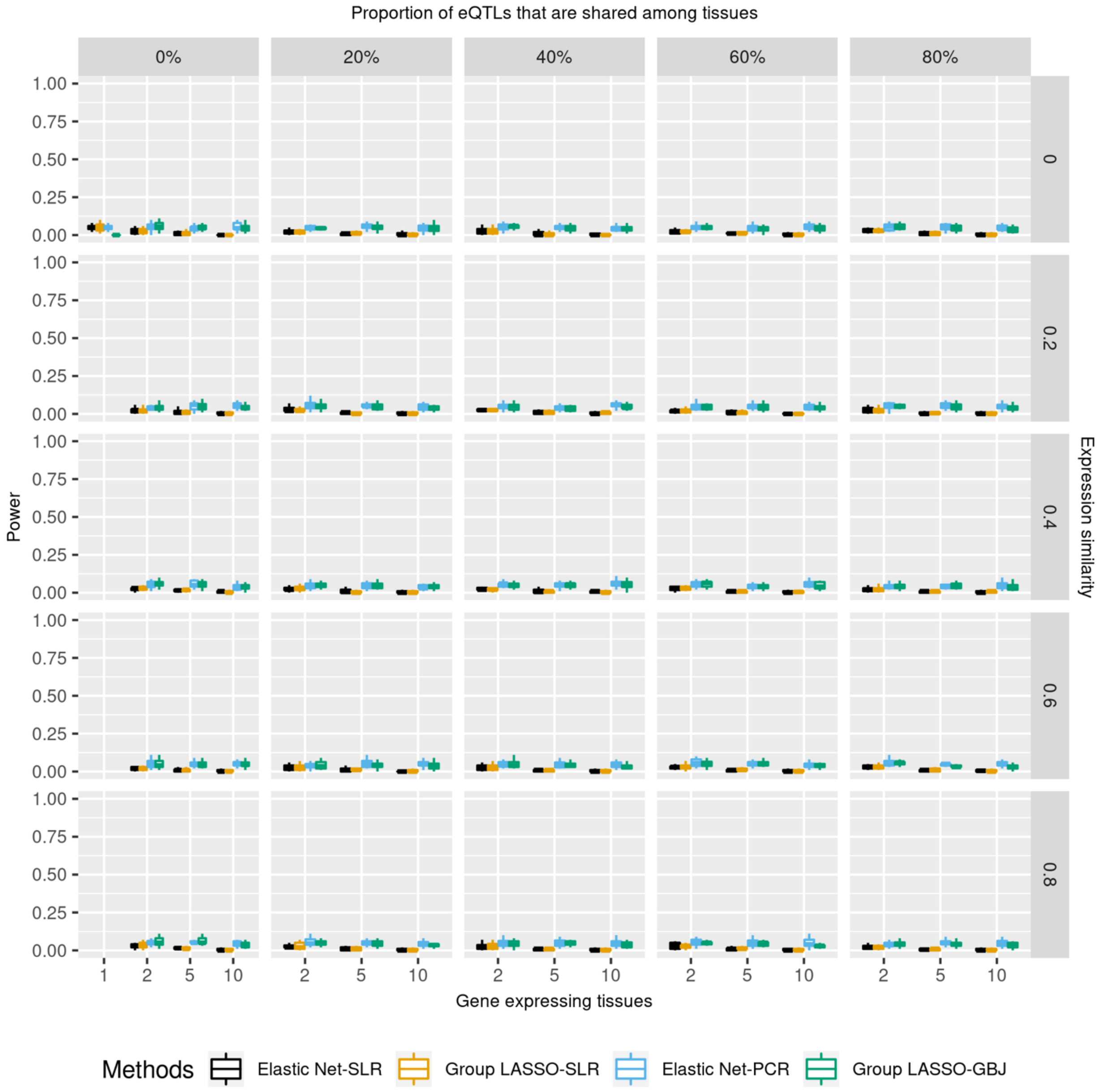


Figure S7. TWAS gene prioritization power when $R^2_{expression-trait}$ = 0.001%

### S11 Fig

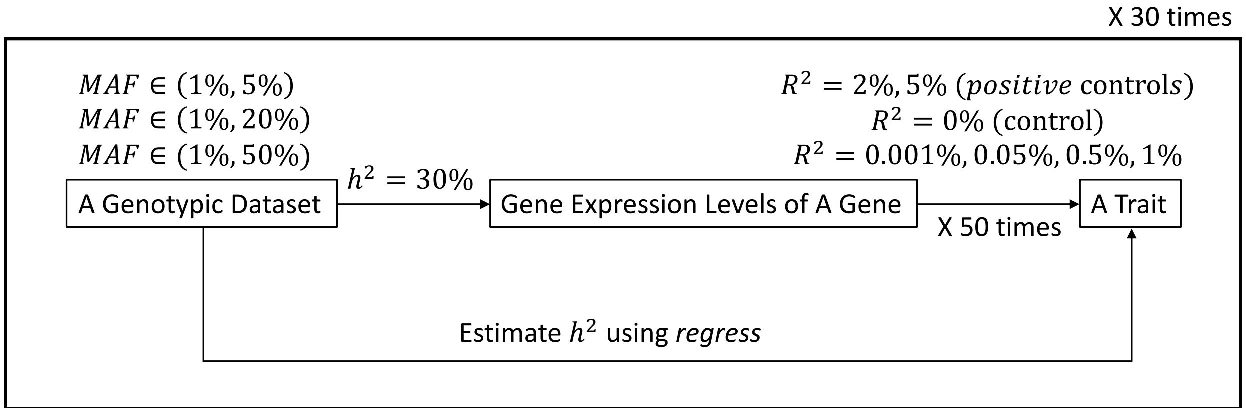


Figure S11. Trait heritability estimation design and workflow

### S12 Fig

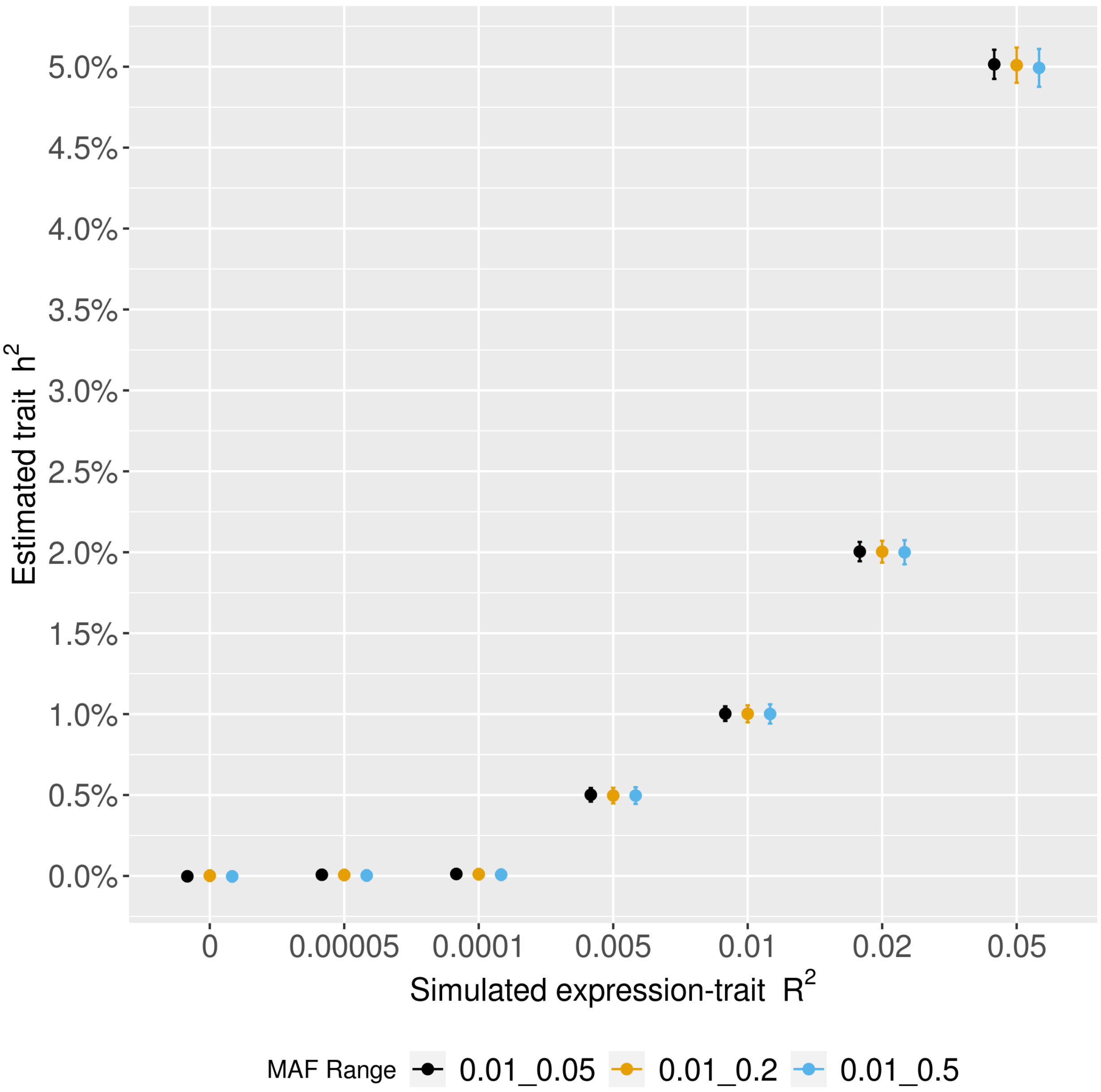


Figure S12. Estimation of trait heritability for simulated datasets.

### S14 Fig

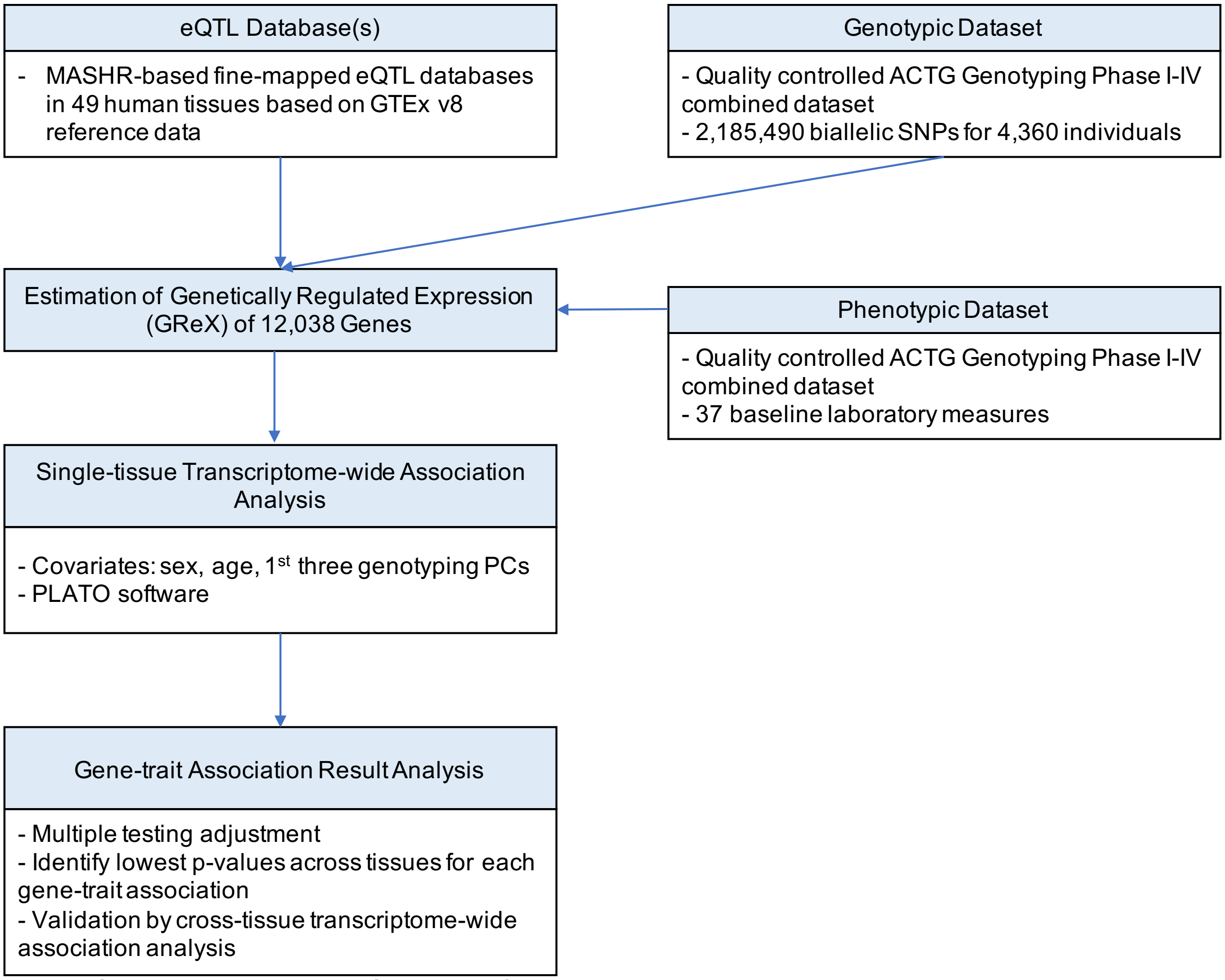

Figure S14. Alternative TWAS analytic framework based on simulation results.
