## Supplementary material for "Tissue specificity-aware TWAS (TSA-TWAS) framework identifies novel associations with metabolic, immunologic, and virologic traits in HIV-positive adults": S8 Fig

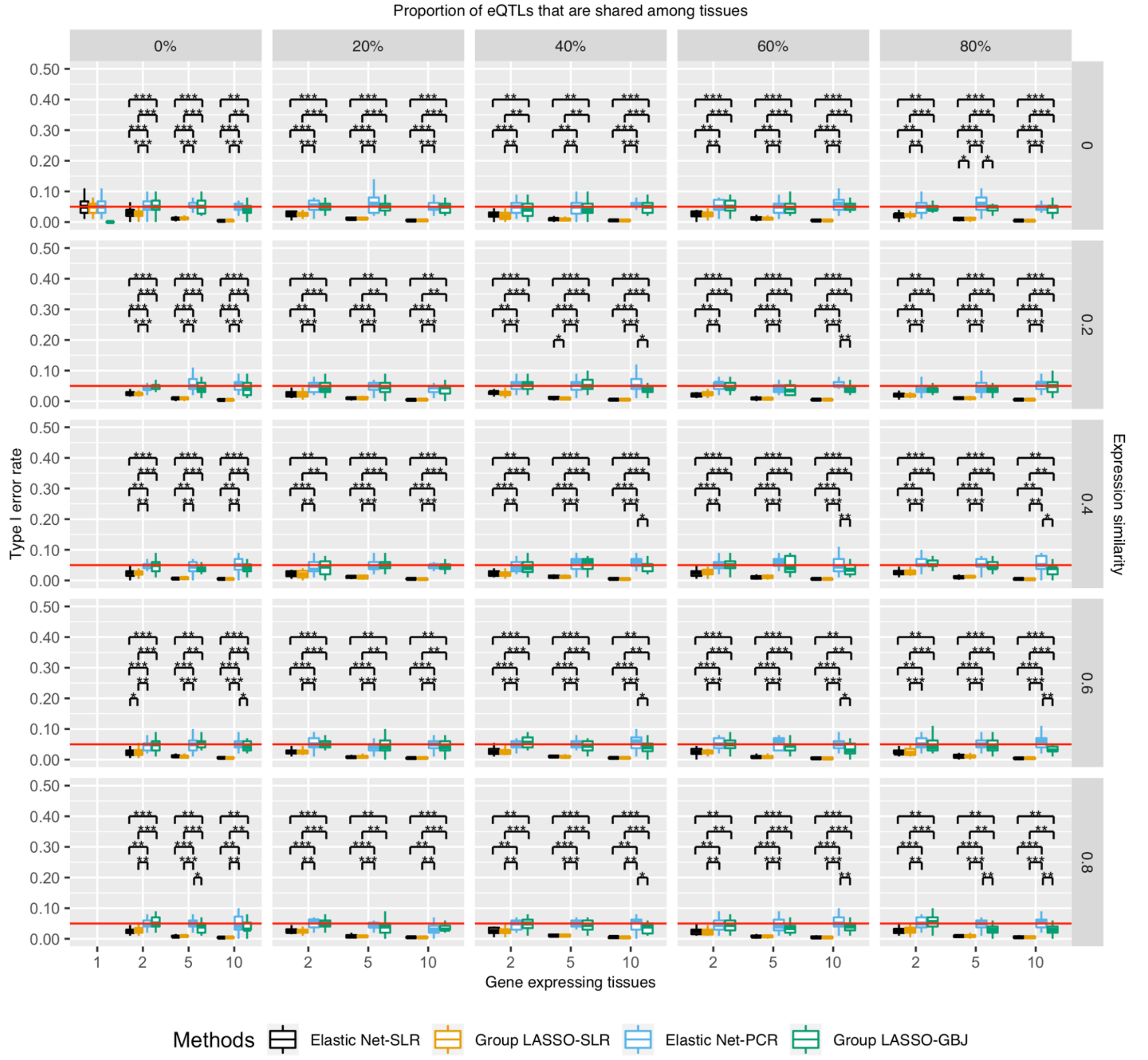


Figure S8. Statistical difference of type I error rates of different TWAS methods. Type I error rate was the probability that TWAS wrongly identified a gene-trait association as significant while there was not any signal. Association p-values were controlled for the number of genes and tested tissues. X-axis is the number of gene-expressing tissues. Each column stands for the proportion of eQTLs that are shared among tissues for a gene. Each row is the similarity of gene expression profiles across tissues which is estimated by correlation. Moving from the top left to the bottom right is a gradient spectrum from tissue-specific genes to broadly expressed genes. The colors represent different TWAS methods and y-axis is the type I error rate. All TWAS methods had controlled type I error rates (⩽ 5%). The difference in type I error rates among different TWAS methods were statistically evaluated (* p-value < 0.05, ** p-value < 0.01, *** p-value < 0.0001).
