## Supplementary material for "Tissue specificity-aware TWAS (TSA-TWAS) framework identifies novel associations with metabolic, immunologic, and virologic traits in HIV-positive adults": S9 Fig

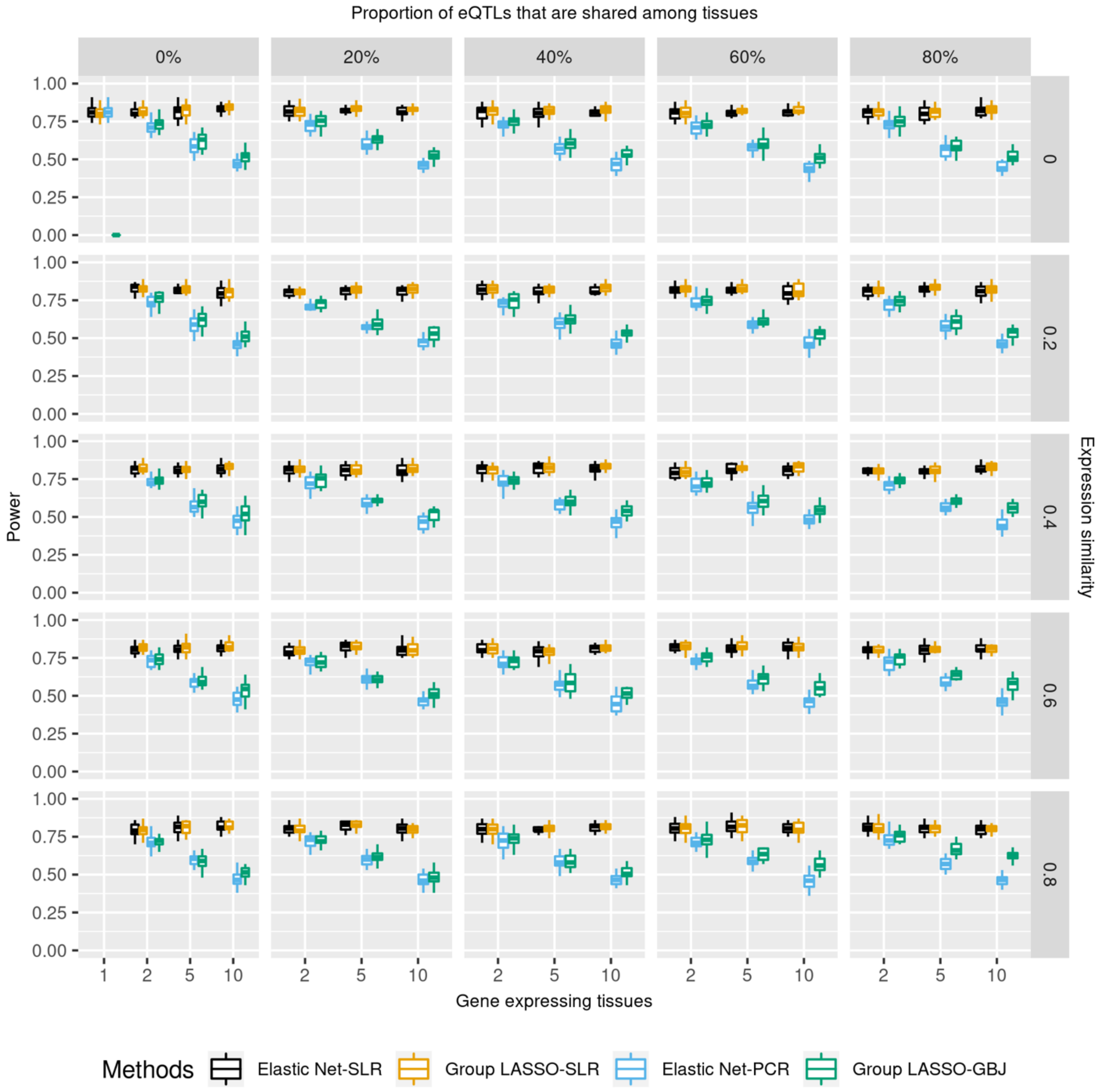


Figure S9. Power when single-tissue associations were not adjusted for the number of tested tissues and $R^2_{expression-trait}$ = 1%.
