## Supplementary material for "Tissue specificity-aware TWAS (TSA-TWAS) framework identifies novel associations with metabolic, immunologic, and virologic traits in HIV-positive adults": S10 Fig

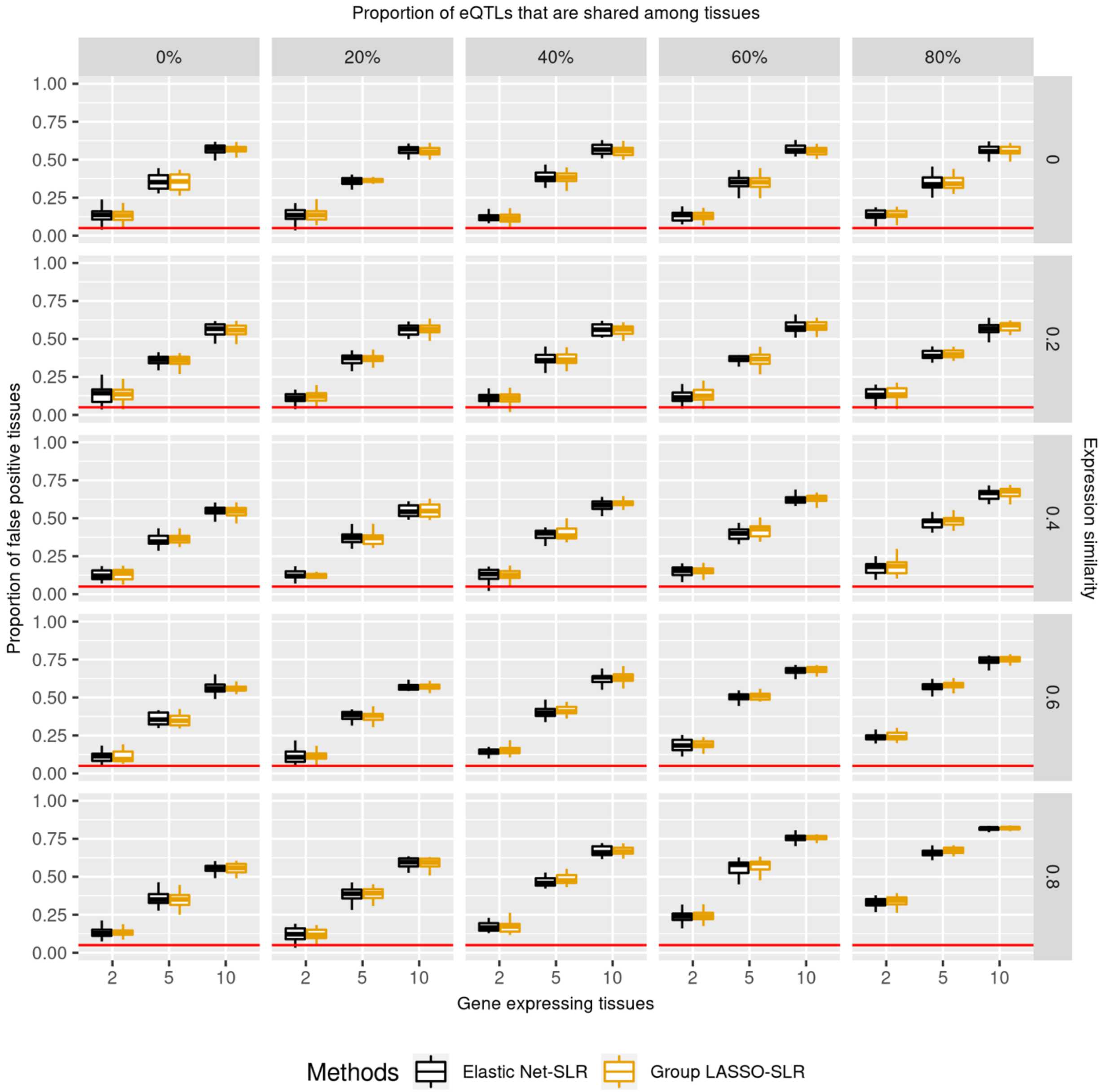


Figure S10. False positive rates when single-tissue associations were not adjusted for the number of tested tissues.
