## Supplementary material for "Tissue specificity-aware TWAS (TSA-TWAS) framework identifies novel associations with metabolic, immunologic, and virologic traits in HIV-positive adults": S13 Fig

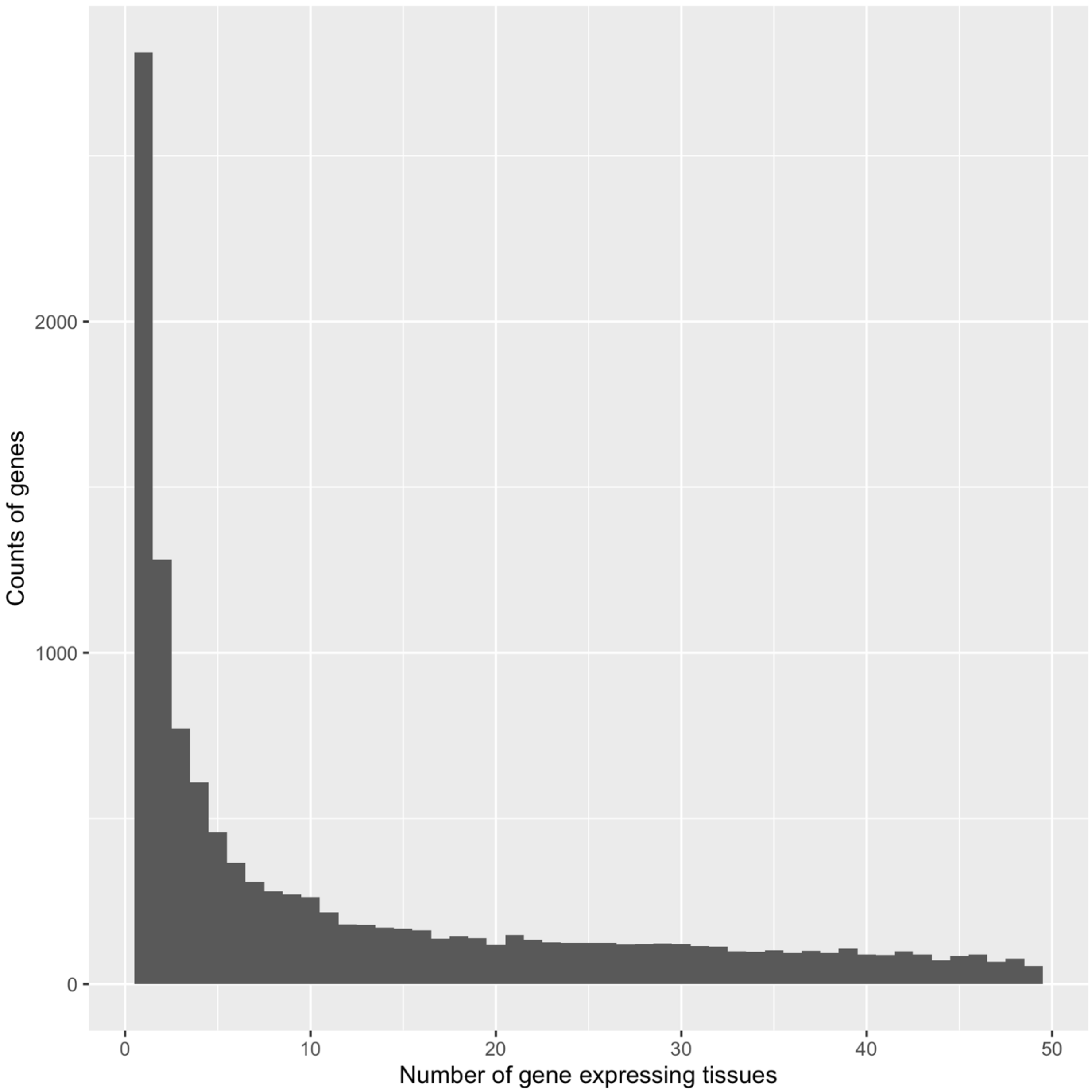


Figure S13. Tissue distribution of predicted GReX. X-axis is the number of tissues that a gene was predicted to be expressed in based on GTEx v8 MASHR-based eQTL models. Y-axis is the count of genes. Genes tended to express in few numbers of tissue according to GTEx v8 prediction.
