## Supplementary material for "Tissue specificity-aware TWAS (TSA-TWAS) framework identifies novel associations with metabolic, immunologic, and virologic traits in HIV-positive adults": S15 Fig

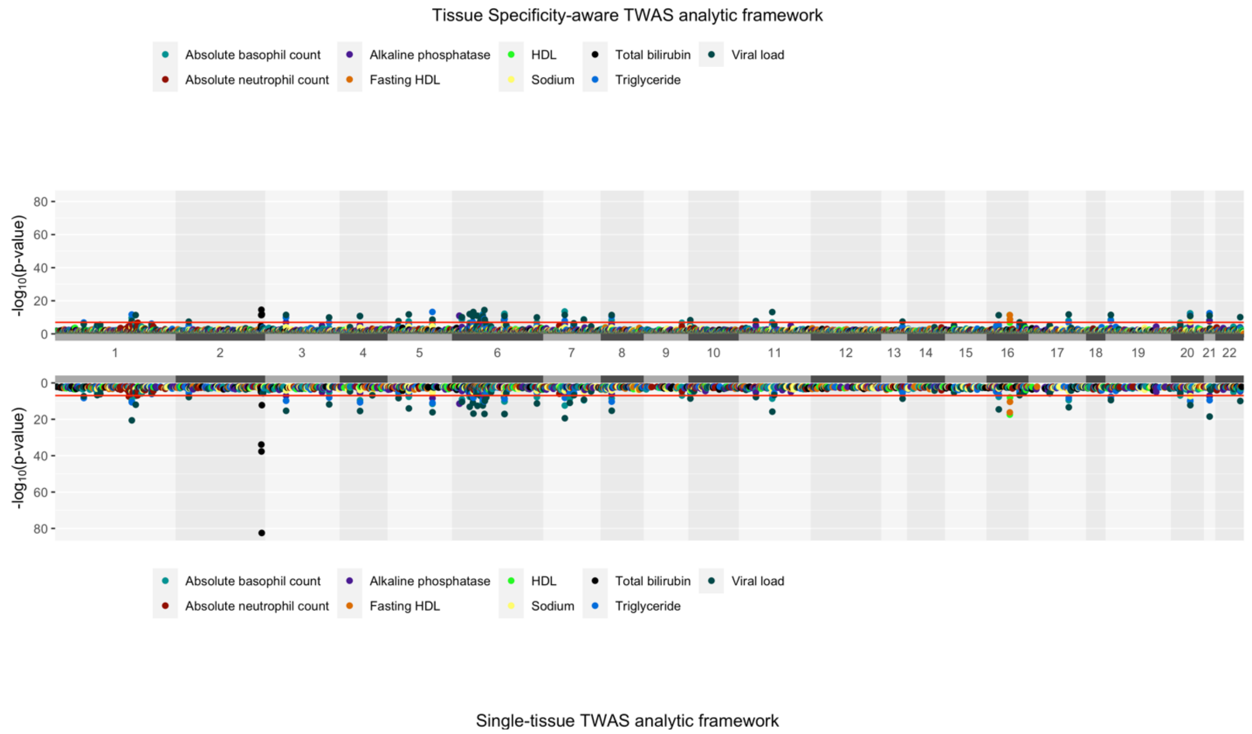


Figure S15. The advantage of our tissue specificity-aware TWAS analytic framework (top) in comparison to a regular single-tissue TWAS. X-axis is the genomic location and y-axis is the -log10 transformed p-values. Colors denote different ACTG baseline laboratories. The significance threshold was $1.12 \times 10^{-7}$ for both top and bottom TWAS frameworks. While single-tissue TWAS were able to find multiple significant gene-trait associations, the significant tissues did not necessarily connect to the traits of interest pathology-wise. Our tissue specificity-aware TWAS framework was able to retain all significant associations that were identified through regular single-tissue TWAS.
