## Supplementary material for "Tissue specificity-aware TWAS (TSA-TWAS) framework identifies novel associations with metabolic, immunologic, and virologic traits in HIV-positive adults": S16 Fig

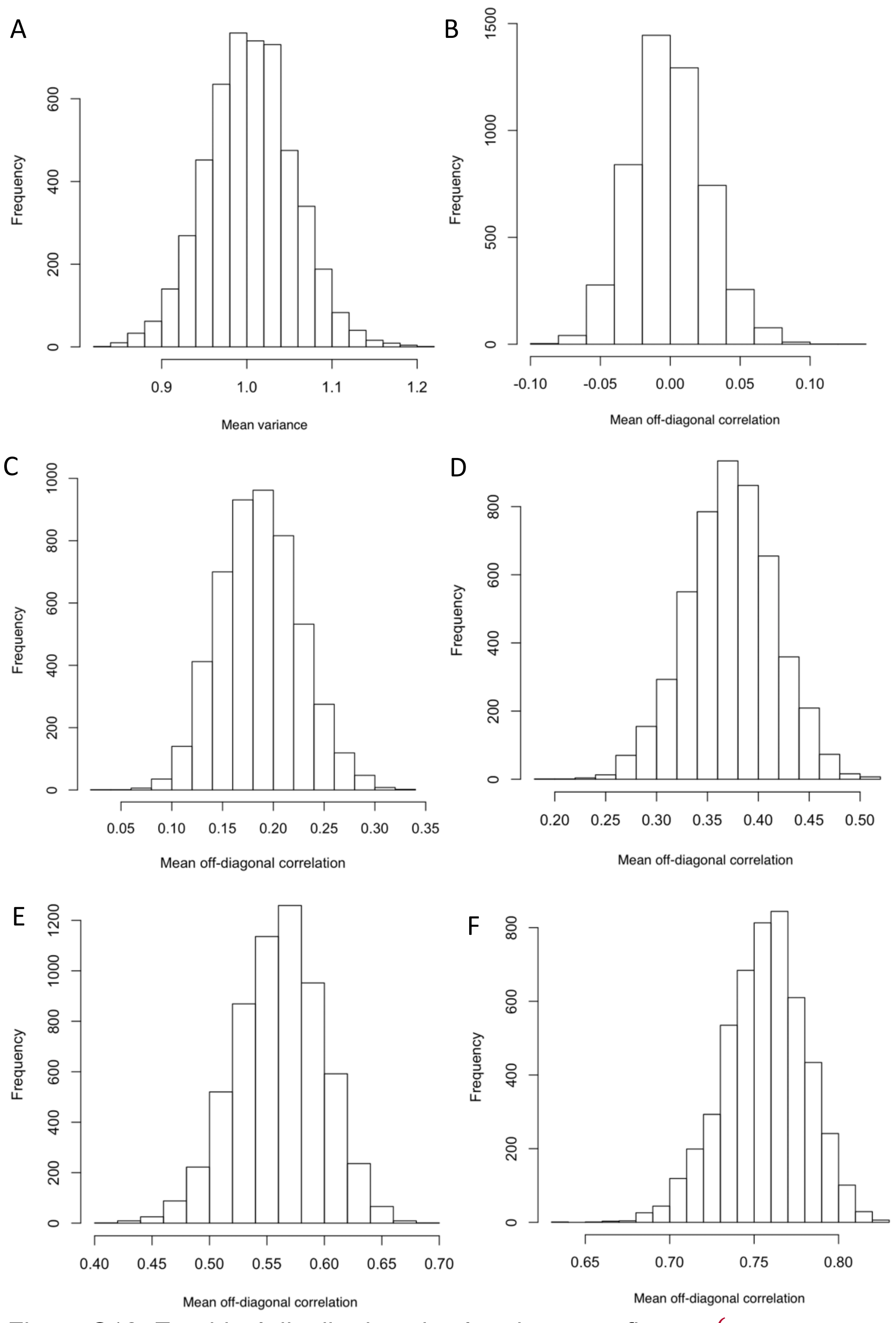


Figure S16. Empirical distribution simulated $E_{N \times P}$ at five $cor(tissue_p, tissue_{p'})$ (0, 0.2, 0.4, 0.6, 0.8). The number of tissues were five in this evaluation. Five thousand rounds of simulations were repeated for each value of $cor(tissue_p, tissue_{p'})$. In each repetition, we obtained one mean variance and one mean off-diagonal correlation across the five simulated tissues. (**A**) The empirical mean variance of $E_{N \times P}$ were approximately one in any situations. The empirical mean off-diagonal correlations euqaled 0, 0.186, 0.373, 0.563, and 0.756, for $cor(tissue_p, tissue_{p'})$ = 0 (**B**), 0.2 (**C**), 0.4(**D**), 0.6 (**E**), 0.8 (**F**).
